# ASCL1 represses a latent osteogenic program in small cell lung cancer in multiple cells of origin

**DOI:** 10.1101/2020.11.11.362632

**Authors:** Rachelle R. Olsen, David W. Kastner, Abbie S. Ireland, Sarah M. Groves, Karine Pozo, Christopher P. Whitney, Matthew R. Guthrie, Sarah J. Wait, Danny Soltero, Benjamin L. Witt, Vito Quaranta, Jane E. Johnson, Trudy G. Oliver

## Abstract

ASCL1 is a neuroendocrine-lineage-specific oncogenic driver of small cell lung cancer (SCLC), highly expressed in a significant fraction of tumors. However, ~25% of human SCLC are ASCL1-low and associated with low-neuroendocrine fate and high MYC expression. Using genetically-engineered mouse models (GEMMs), we show that alterations in *Rb1/Trp53/Myc* in the mouse lung induce an ASCL1^+^ state of SCLC in multiple cells of origin. Genetic depletion of ASCL1 in MYC-driven SCLC dramatically inhibits tumor initiation and progression to the NEUROD1^+^ subtype of SCLC. Surprisingly, ASCL1 loss converts tumors to a SOX9^+^ mesenchymal/neural-crest-stem-like state that can differentiate into RUNX2^+^ bone tumors. ASCL1 represses SOX9 expression, as well as WNT and NOTCH developmental pathways, consistent with human gene expression data. Together, SCLC demonstrates remarkable cell fate plasticity with ASCL1 repressing the emergence of non-endodermal stem-like fates that have the capacity for bone differentiation.

---

Small cell lung cancer (SCLC) is a highly aggressive neuroendocrine lung tumor with a median survival time of only 10-12 months^1,2^. Almost all SCLCs exhibit inactivation of the tumor suppressor genes *RB1* and *TP53*, along with mutually exclusive gain of a MYC family member including *MYC, MYCL* or *MYCN*^3–5^. While SCLC has historically been treated as a single disease in the clinic, it is increasingly appreciated to be composed of distinct molecular subtypes^1^. Approximately 70% of SCLCs are characterized by high expression of the transcription factors *ASCL1* and *MYCL*, along with high expression of neuroendocrine (NE) markers such as *Synaptophysin* (*SYP*), *Ubiquitin C-terminal Hydrolase L1* (*UCHL1*) and *Chromogranin A* (*CHGA*)^1,4–6^. NE-high SCLCs typically display classic morphology with small round blue cells and scant cytoplasm. In contrast, approximately 25% of SCLCs are associated with an ASCL1-low/NE-low gene expression profile, and these tend to have high *MYC* expression and variant morphology. We reported the first genetically-engineered mouse model (GEMM) of MYC-driven SCLC^6^, and demonstrated that MYC promotes the variant, ASCL1-low/NE-low subtype of SCLC. Importantly, the MYC-driven subtype of SCLC demonstrates unique therapeutic vulnerabilities including sensitivity to inhibition of Aurora kinases (AURK), CHK1, IMPDH, and arginine depletion^6–11^.

ASCL1 and NEUROD1 are lineage-specifying basic helix-loop-helix transcription factors that activate neuroendocrine genes and are required for neural differentiation^1,12^. Genetic knockout studies in the mouse demonstrate that ASCL1 is essential for the development of pulmonary neuroendocrine cells (PNECs)^13^, and ASCL1 is expressed in the adult lung specifically in PNECs^14^. Importantly, ASCL1 is necessary for the development of classic SCLC as demonstrated in the *Rb1^fl/fl^;Trp53^fl/fl^;Rbl2^fl/fl^* (RPR2) mouse model where conditional *Ascl1* deletion abolishes tumor formation^12^. NEUROD1 is not required for SCLC initiation or progression in the RPR2 model, consistent with the observation that NEUROD1 is not expressed in classic SCLC models^6,12^. NEUROD1 is expressed in the variant MYC-driven SCLC model, and is associated with MYC expression in human SCLC^6^. NEUROD1 has recently been suggested to temporally follow ASCL1 during MYC-driven subtype evolution^15^. However, the function of ASCL1 and NEUROD1 in MYC-driven tumors in vivo has not yet been determined.

PNECs have been generally accepted to be the cell of origin for SCLC, although recent studies have challenged whether they are the only relevant initiating cell population^8,15–17^. SCLC shares molecular similarities with normal PNECs, including expression of ASCL1 and its neuroendocrine target genes. Studies in mouse models have demonstrated that neuroendocrine cells are the most permissive for SCLC development in the original classic *Rb1^fl/fl^;Trp53^fl/fl^* (RP) model and in RPR2 mice^18,19^. Club cells and alveolar type II (AT2) cells in these studies were relatively resistant to SCLC transformation^18,19^. More recently, a subset of SCLC was revealed to harbor a tuft-cell-like signature with dependency on the tuft-cell transcription factor POU2F3^8^, suggesting that the tuft cell may serve as a cell of origin for SCLC. Basal cells have also been described as cells of origin for SCLC^16,20^. Together, these studies suggest that lung cells may have broad plasticity for SCLC transformation, but the cells of origin for MYC-driven SCLC have not been determined.

Here, we use new GEM models to determine the function of ASCL1 in MYC-driven SCLC derived from multiple cells of origin.

## Results

### MYC-driven SCLC can arise in multiple lung cell types

Our previous work demonstrated that *Myc* overexpression in the *Rb1^fl/fl^;Trp53^fl/fl^;MycT58A^LSL/LSL^* (RPM) mouse model promotes the development of a variant, neuroendocrine-low subtype of SCLC that recapitulates features of the human disease^6^. To determine which cell types in the normal lung epithelium are capable of initiating tumorigenesis in the RPM model, we intratracheally-infected mice with adenoviruses carrying cell-type-specific promoters driving expression of *Cre recombinase*. We used a general *Cmv* promoter, a neuroendocrine *Cgrp* promoter, a club cell *Ccsp* promoter, and an AT2 *Spc* promoter, whose specificity have been previously described^17–19,21^. The combination of *Myc* overexpression with *Rb1* and *Trp53* loss was sufficient to initiate tumorigenesis with all four Cre viruses (Fig. 1a and 1b). RPM-CMV tumors had the shortest latency (43 days median survival), followed by RPM-CGRP tumors (55 days median survival). Compared to results from the RP model^18^, targeting of club cells with CCSP-Cre led to tumors with a surprisingly short latency (77 days median survival), and was almost as efficient at tumor initiation as targeting neuroendocrine cells, suggesting that club cells are highly susceptible to MYC-mediated SCLC transformation. Consistent with the location of club cells, in situ tumors in the CCSP-Cre RPM mice were often located within bronchioles (Supplementary Fig. 1a). In contrast, AT2 cells were the most resistant to transformation as RPM tumors initiated with SPC-Cre had the longest latency (184 days median survival) (Fig. 1a). Review by board-certified pathologists (A. Gazdar and B. Witt) confirmed that RPM mice infected with different cell-type-specific Cre viruses predominantly developed SCLC, with a mixture of classic and variant histopathologies and perilymphatic spread, similar to that observed in the original RPM-CGRP model (Fig. 1b and Supplementary Fig. 1a). Combined with previous studies in the RP and RPR2 mice^17–19^, these results suggest that genetic alterations can dictate the susceptibility of cells to serve as a cell of origin for SCLC.

**Figure 1:**
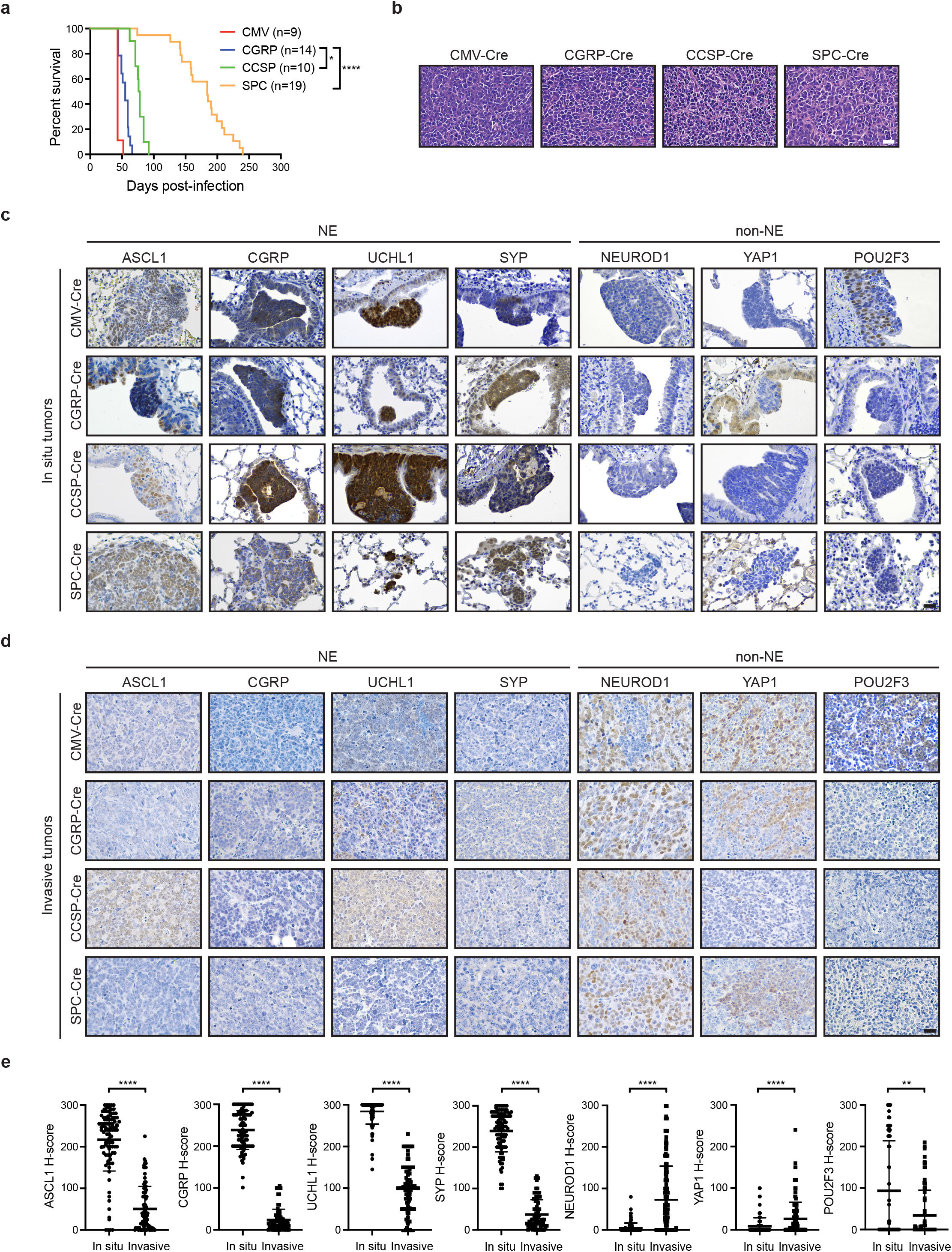
MYC-driven SCLC can arise in multiple lung cell types and initially expresses ASCL1. a) Survival of RPM mice infected with indicated cell-type-specific Ad-Cre viruses. Number of mice indicated in the figure. Mantel-Cox log-rank test, * p < 0.05; **** p < 0.0001. b) Representative H&E histology of SCLC from RPM mice initiated with indicated Ad-Cre viruses. Scale bar: 25 μm. c) Representative immunohistochemistry (IHC) of early in situ tumor lesions for indicated antibodies (top) in RPM mice infected with indicated Ad-Cre viruses (left). Scale bar: 25 μm. d) Representative IHC of large, invasive tumors for indicated antibodies (top) in RPM mice infected with indicated Ad-Cre viruses (left). Scale bar: 25 μm. e) H-score quantification of IHC in panels (c) and (d). H-Score = % of positive cells multiplied by intensity score of 0-3; see Methods. Approximately 70-200 tumors from n ≥ 3 mice per condition were quantified. Data is shown as mean ± standard deviation (SD). Mann-Whitney two-tailed t-tests, **** p < 0.0001; ** p < 0.01. See also Supplementary Figure 1.

### MYC-driven SCLC induces ASCL1 in multiple cells of origin

To further determine whether tumors in this model were neuroendocrine, we examined in situ and invasive tumors for expression of ASCL1 and other neuroendocrine markers including CGRP, UCHL1 and SYP. Interestingly, in situ tumors from each cohort were dominated by an ASCL1^high^/neuroendocrine^high^ (NE^high^) phenotype even when tumors were initiated in non-neuroendocrine cells with CCSP-Cre or SPC-Cre (Fig. 1c). NE^high^ in situ tumors did not express CCSP or SPC (Supplementary Fig. 1b), suggesting that the original cell fate identities were rapidly lost. Consistent with the classic NE^high^ state of these in situ lesions, the majority of in situ RPM tumors expressed low-to-undetectable levels of NEUROD1, YAP1, and POU2F3 (Fig. 1c), key transcription factors that are associated with NE^low^ subtypes of SCLC^1^. These results suggest that MYC overexpression in the context of *Rb1/Trp53* loss alters the original cellular differentiation state and initially reprograms cells toward a neuroendocrine-like progenitor.

In contrast to the in situ tumors, invasive RPM tumors were dominated by an ASCL1^low^/NE^low^ phenotype largely independent of the cell of origin, and tended to express higher levels of NEUROD1 and YAP1, but not POU2F3 (Fig. 1d and 1e). Interestingly, tumors initiated in club cells exhibited significantly higher ASCL1 and less YAP1 (Supplementary Fig. 1c), suggesting club cells may not progress as readily as tumors from other cells of origin. Although POU2F3 levels were not significantly elevated in all of the models, we observed an increase in POU2F3^+^ invasive tumors specifically in the RPM-CMV mice (Supplementary Fig. 1c), consistent with recent studies^15^ suggesting that the *Cmv* promoter may occasionally be targeting a POU2F3^+^ tuft cell^8^. Together, this suggests that RPM tumors induce ASCL1 and a neuroendocrine phenotype in multiple cells of origin that progress to a NE-low state.

### ASCL1 loss delays tumorigenesis and promotes bone differentiation in multiple cells of origin

Since ASCL1 has been previously shown to be required for development of classic SCLC in the RPR2 model^12^, we sought to determine the role of ASCL1 in MYC-driven variant SCLC. We crossed RPM mice to *Ascl1*-floxed animals to generate *Rb1^fl/fl^;Trp53^fl/fl^;MycT58A^LSL/LSL^;Ascl1^fl/fl^* (RPMA) mice and infected RPMA mice with cell type-specific adenoviruses as described above. In contrast to prior findings in classic SCLC models^12^, RPMA mice developed tumors in the lung, albeit with significantly delayed latencies compared to RPM mice (Fig. 2a). RPMA-CMV mice developed tumors with the shortest median survival of 81 days (1.9-fold longer than RPM-CMV mice), while RPMA mice infected with CGRP-Cre had a median survival of 139 days (2.5-fold longer than RPM-CGRP mice). Median survival in the RPMA-CCSP mice was 256 days (3.3-fold longer than RPM-CCSP mice). RPMA-SPC tumors had the longest latency (median survival undetermined; at least 2.4-fold longer than RPM-SPC mice). Successful Cre-mediated recombination of the *Rb1, Trp53*, and *Ascl1* alleles in these mice was assessed by genomic PCR on individual micro-dissected RPMA tumors (Supplementary Fig. 2a). We observed recombined *Rb1*, *Trp53* and *Ascl1* alleles in ~94% of tumors. Unrecombined alleles were also detected in the majority of samples, but we cannot rule out that this represents normal tissue contamination in each tumor since it is difficult to cleanly dissect these tumors. We also detected rare tumors that did not recombine the *Ascl1* allele, and in these cases, the tumors expressed ASCL1 protein and exhibited SCLC morphology (Supplementary Fig. 2b).

**Figure 2:**
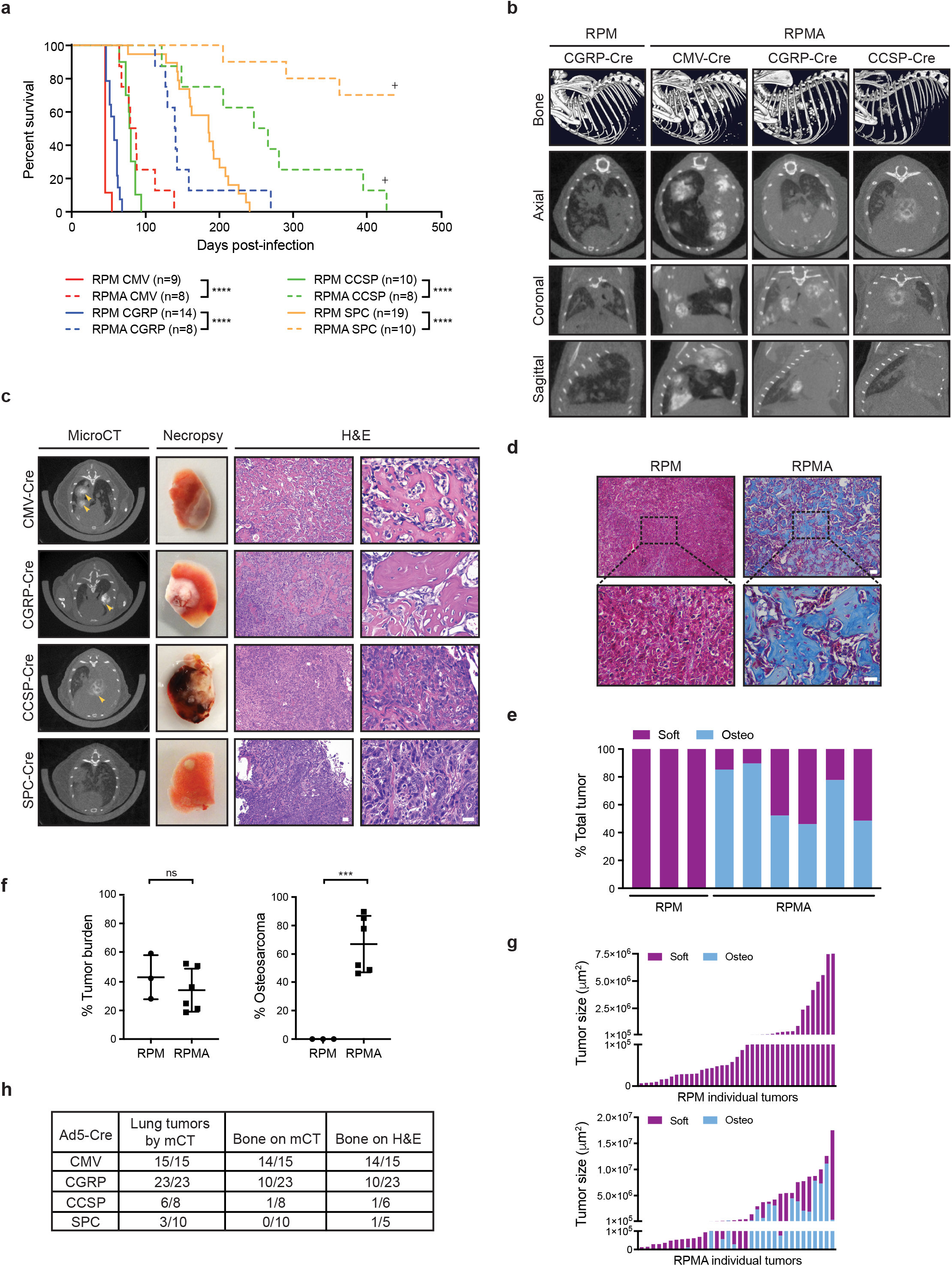
ASCL1 loss delays tumorigenesis and promotes bone differentiation in multiple cells of origin. a) Survival curve comparing RPM (solid lines, data from Figure 1a) vs RPMA mice (dashed lines) infected with indicated cell-type-specific Cre viruses. Number of mice indicated in the figure. + = mouse censored to end the cohort. Mantel-Cox log-rank test, **** p < 0.0001. b) Representative bone analysis (top row) of microCT images in RPM vs RPMA mice with advanced lung tumors infected with the indicated Ad-Cre viruses. Matched axial, coronal, and sagittal cross-sections of RPM and RPMA tumors used for bone analysis (bottom rows). c) Representative microCT axial cross-sections, necropsy images, and H&E staining for tumors from mice infected with indicated Ad-Cre viruses. In microCT panel, yellow arrowheads indicate areas of focal bone formation. Necropsy image shows one whole lung lobe. In H&E panels, scale bar indicates 50 μm in 10x image (left), and 25 μm in the 40x image (right). d) Representative Trichrome staining in RPM-CMV vs RPMA-CMV tumors. Scale bar indicates 50 μm in the 10x image (top) and 25 μm in the 40x image (bottom). e) Quantification of the percent soft vs osteosarcoma tumor in RPM-CMV (n=3) vs RPMA-CMV (n=6) mice with equivalent total tumor burden determined by PAS with Alcian Blue (PAB) staining. f) Average percent total tumor burden and total osteosarcoma in RPM vs RPMA mice derived from Figure 2e. Mean ± SD. Unpaired two-tailed t-test, *** p < 0.001; ns = not significant. g) Size of individual soft tumors and osteosarcomas quantified from PAB-stained slides from a representative RPM-CMV (top) or RPMA-CMV (bottom) mouse. h) Table indicating number of RPMA mice per cohort with lung tumors detected by microCT imaging, bone detected by microCT, or bone detected by H&E review. See also Supplementary Figure 2 and Supplementary Videos.

While imaging RPMA mice by microCT, we made the unexpected observation that mice were developing lung tumors with a tissue density consistent with bone (Fig. 2b). We used bone analysis imaging software on the Quantum GX2 microCT to compare lung tumor development in RPM and RPMA mice. Bone analysis of RPM mice with high lung tumor burden identified only the skeleton (Fig. 2b, top panel, and Supplementary Videos). In contrast, bone analysis of RPMA-CMV, -CGRP, and -CCSP animals identified both the skeleton as well as multiple bone-like tumors within the lung area. These lesions were clearly evident on multiple microCT cross-sections (Fig. 2b, bottom panels, and Supplementary Videos). At necropsy, many RPMA lesions were mineralized and required decalcification prior to sectioning (Fig. 2c). Analysis of H&E-stained tissues by a board-certified pathologist (B.W.) confirmed that RPMA lungs contained high-grade osteosarcoma with well-developed osteoid (Fig. 2c, right panels, and Supplementary Fig. 2c). Bone-like lesions in RPMA tumors were also confirmed by Trichrome staining (Fig. 2d), Periodic Acid Schiff (PAS) with Alcian Blue staining (PAB, Supplementary Fig. 2d), and Toluidine blue staining (Supplementary Fig. 2e). Overall, osteosarcomas represented between 45-90% of the total tumor burden in RPMA-CMV animals compared to RPM animals that had no osteosarcomas detected despite equivalent tumor burden (Fig. 2e and 2f). Small RPMA tumors were primarily soft, while large invasive RPMA tumors displayed a mix of soft and osteosarcoma phenotypes (Fig. 2g), suggesting that tumors may arise in a dedifferentiated state that later adopts a bone phenotype. We verified that the vast majority of both in situ and invasive RPMA tumors lacked ASCL1 expression (Supplementary Fig. 2f). Some RPMA mice had lymph node metastases, including one animal that appeared to have an osteosarcoma metastasis (Supplementary Fig. 2g). RPMA mice also had occasional non-calcified tumors resembling adenocarcinoma or large cell carcinoma that lacked ASCL1 expression (Supplementary Fig. 2h). The osteosarcoma-like tumors were frequently observed by microCT imaging in RPMA-CMV and RPMA-CGRP mice, less frequent in RPMA-CCSP mice, and not detectable in RPMA-SPC mice (Fig. 2h). However, one RPMA-SPC mouse did have areas of focal bone formation with early osteoid detected upon histopathological review (Fig. 2h). These data suggest that in the context of MYC-driven SCLC, ASCL1 represses a latent osteogenic fate in multiple lung cell types that may have differing propensities for bone differentiation.

### RPMA tumors are transcriptionally distinct with loss of ASCL1 and NEUROD1 target genes

Following the striking observation that loss of ASCL1 in MYC-driven SCLC promotes osteosarcoma formation, we sought to better understand the transcriptional program of RPMA tumors. We performed RNA-sequencing (RNA-seq) on variant RPM (n = 11), classic RPR2 (n = 6), and un-calcified RPMA tumors (n = 7) including one RPMA tumor that was heterozygous for *Ascl1 (Rb1^fl/fl^;Trp53^fl/fl^;MycT58A^LSL/LSL^;Ascl1^fl/wt^; RPMA*^het^). Consistent with their histological differences, principal component analysis of RNA-seq data revealed distinct clustering of all three tumor types (Fig. 3a). Notably, the one heterozygous RPMA^het^ tumor clustered apart from homozygous *Ascl1*-null RPMA samples and closer in identity to the RPR2 and RPM samples. Thus, complete loss of *Ascl1* appears to be required for the osteoid gene expression signature, consistent with histological observations.

**Figure 3:**
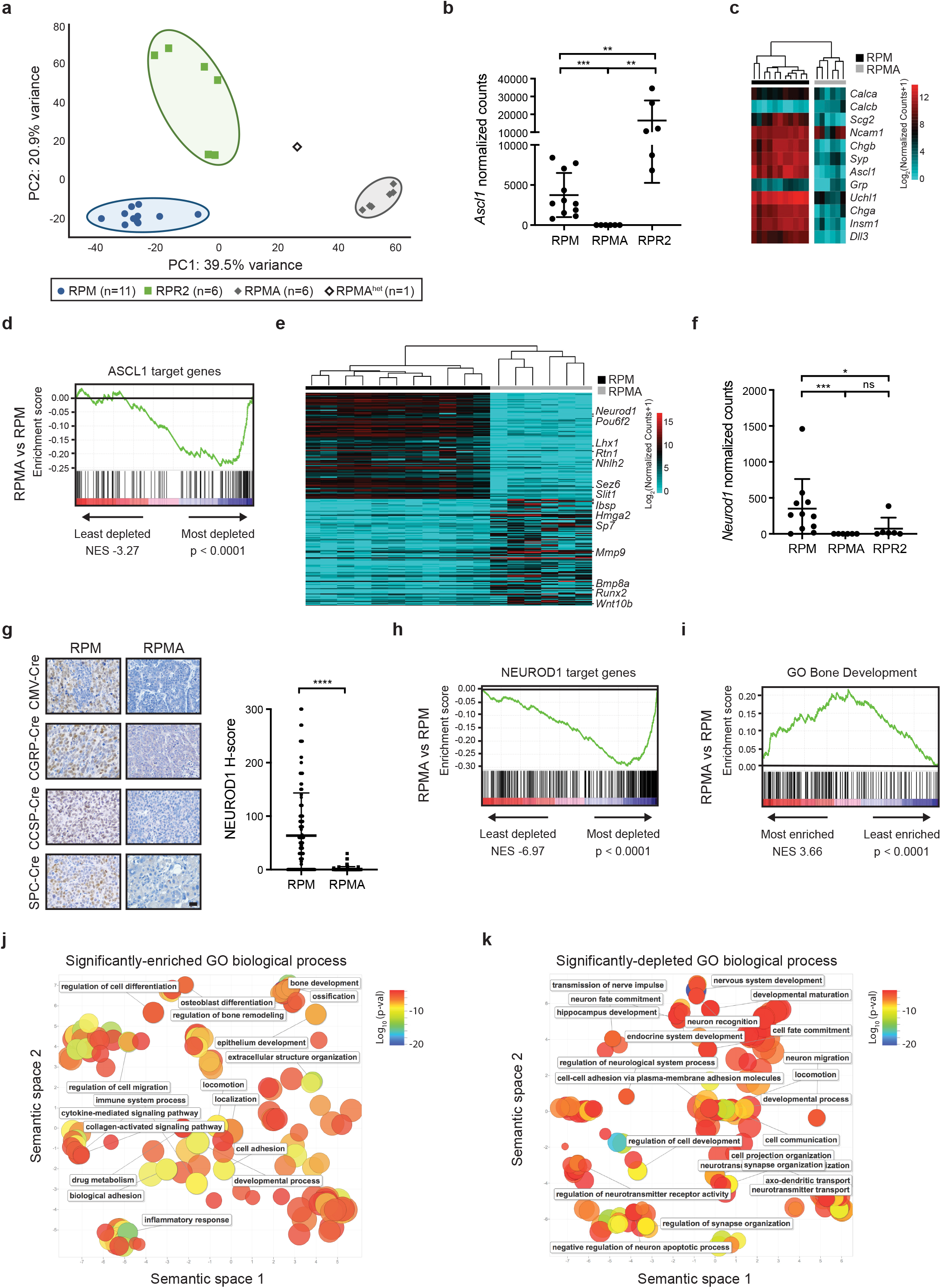
RPMA tumors are transcriptionally distinct with loss of ASCL1 and NEUROD1 target genes. a) Principal component (PC) analysis comparing gene expression by bulk RNA-seq in RPM, RPMA, RPMA^het^ (*Rb1^fl/fl^;Trp53^fl/fl^;Myc^LSL/LSL^;Ascl1^fl/wt^*), and RPR2 tumors. b) *Ascl1* expression shown as normalized counts by RNA-seq from lung tumors in indicated GEMMs. Mean ± SD. Mann-Whitney two-tailed t-test, ** p < 0.01; *** p < 0.0003. c) Heat map comparing log2 normalized counts for expression of select neuroendocrine genes in RPM vs RPMA tumors. d) Gene set enrichment analysis (GSEA) in RPMA vs RPM tumors using ASCL1 ChIP-seq target genes from Borromeo et al, *Cell Rep*, 2016. Normalized enrichment score (NES) and p value indicated in figure. e) Heat map of top 200 (100-up and 100-down) differentially-expressed genes in RPMA vs RPM tumors with select genes indicated. f) *Neurod1* expression as normalized counts by RNA-seq from lung tumors in indicated GEMMs. Mean +/- SD. Mann-Whitney two-tailed t-test, * p < 0.05; *** p < 0.001; ns = not significant. g) Representative IHC and H-score quantification for NEUROD1 in RPM vs RPMA tumors. Approximately 18-80 tumors were quantified from n ≥ 4 mice per condition. Scale bar: 25 μm. Mean +/- SD. Mann-Whitney two-tailed t-test, **** p < 0.0001. h) GSEA comparing RPMA vs RPM tumors using NEUROD1 ChIP-seq target genes from RPM tumors and human SCLC cell lines from Borromeo et al, *Cell Rep*, 2016. Normalized enrichment score (NES) and p value indicated in the figure. i) GSEA comparing RPMA vs RPM tumor gene expression to a known ossification signature, “GO_Bone_Development.” NES and p values indicated in the figure. j) Scatterplot visualizing semantic similarity of GO Biological Processes enriched in RPMA vs RPM tumors. k) Scatterplot visualizing semantic similarity of GO Biological Processes depleted in RPMA vs RPM tumors.

We verified that *Ascl1* mRNA was significantly depleted in RPMA tumors (Fig. 3b). Consistently, established ASCL1 target genes including neuroendocrine genes were significantly reduced in RPMA compared to RPM tumors (Fig. 3c and 3d). We next performed differential gene expression analysis to identify additional genes and pathways that discriminate the RPMA and RPM tumors. Interestingly, these data revealed *Neurod1* as one of the most significantly down-regulated genes in RPMA versus RPM tumors (Fig. 3e) with complete loss of *Neurod1* transcript counts (Fig. 3f). As assessed by IHC, RPMA tumors were devoid of NEUROD1 protein, in contrast to high levels observed in RPM tumors (Fig. 3g). Gene-set enrichment analysis (GSEA) using established human NEUROD1 target genes^12^ combined with NEUROD1 ChIP-seq from mouse RPM tumors (n = 2) revealed a significant depletion of NEUROD1 target genes in RPMA compared to RPM tumors (Fig. 3h). These data reveal that deletion of ASCL1 in RPM tumors during tumor initiation abolishes the NEUROD1^+^ subtype of SCLC, consistent with recent findings that *Ascl1* expression temporally precedes *Neurod1* expression during MYC-driven SCLC progression^15^.

Consistent with the observed osteoid formation, GSEA revealed a significant positive enrichment for bone development genes in RPMA compared to RPM tumors (Fig. 3i). To identify other biological processes altered following *Ascl1* loss in RPM mice, we performed GO Enrichment Analysis on all significantly enriched or depleted genes in RPMA compared to RPM tumors (Fig. 3j and 3k). As expected, ossification-related processes were significantly upregulated in RPMA, as well as immune-related signatures (Fig. 3j). Significantly-depleted biological processes in RPMA tumors included those related to neuronal development (Fig. 3k), consistent with the loss of ASCL1 and NEUROD1 in these tumors. Together, these data highlight a critical role for ASCL1 in promoting neuroendocrine cell fate and repressing an underlying osteosarcoma-like fate in MYC-driven SCLC.

### Network analyses predict transcriptional regulators that drive osteosarcoma cell fate upon ASCL1 loss

To identify the key transcription factors that are responsible for this dramatic change in cell fate upon ASCL1 loss, we turned to Weighted Gene Coexpression Network Analysis (WGCNA). First, we generated a coexpression network of all genes, which allowed the identification of distinct gene modules across all RPM and RPMA samples. Using an ANOVA statistical test, we identified the gene modules that significantly differed between RPM and RPMA tumors (Fig. 4a and Supplementary Fig. 3a). Strikingly, ~1/3 of the transcriptome was altered upon ASCL1 loss. To generate a gene regulatory network, we focused on transcription factors predicted by WGCNA to be central to differentially-expressed gene modules, as well as known regulators of lung cancer cell fate. Using BooleaBayes^22^, we determined rules of interaction between these transcription factors, where the rules underlie the dynamics of cell identity in the presence or absence of ASCL1 (Supplementary Fig. 3b). For example, ASCL1 is regulated by eight parent nodes (AR, E2F1, HES1, KLF4, MITF, NR3C1, PHC1, and RUNX1). Each ON/OFF combination of these parent nodes determine the expression of ASCL1. Likewise, ASCL1 regulates expression of a number of downstream transcription factors. These regulations define how a cell may change its identity or reach a stable phenotype (an attractor state). Dynamic simulations identified two attractor states, each corresponding to either RPM or RPMA tumors (Fig. 4b). As described in [22], we used a random walk simulation to predict regulators driving these steady states. Satisfyingly, *in silico* silencing of the ASCL1 or NEUROD1 node destabilized the RPM attractor, consistent with experimental results (Fig. 4b). Conversely, activation of ASCL1 or NEUROD1 in the RPMA attractor destabilized that steady state. This is reminiscent of human SCLC, in which ASCL1 is a destabilizer of the non-neuroendocrine subtype (SCLC-Y)^22^. By comparing human SCLC cell lines to mouse tumors using PCA analysis (Fig. 4c), RPMA tumors clustered with the non-neuroendocrine POU2F3 and YAP1 subtypes, suggesting a similarity between RPMA tumors and human non-neuroendocrine subtypes.

**Figure 4:**
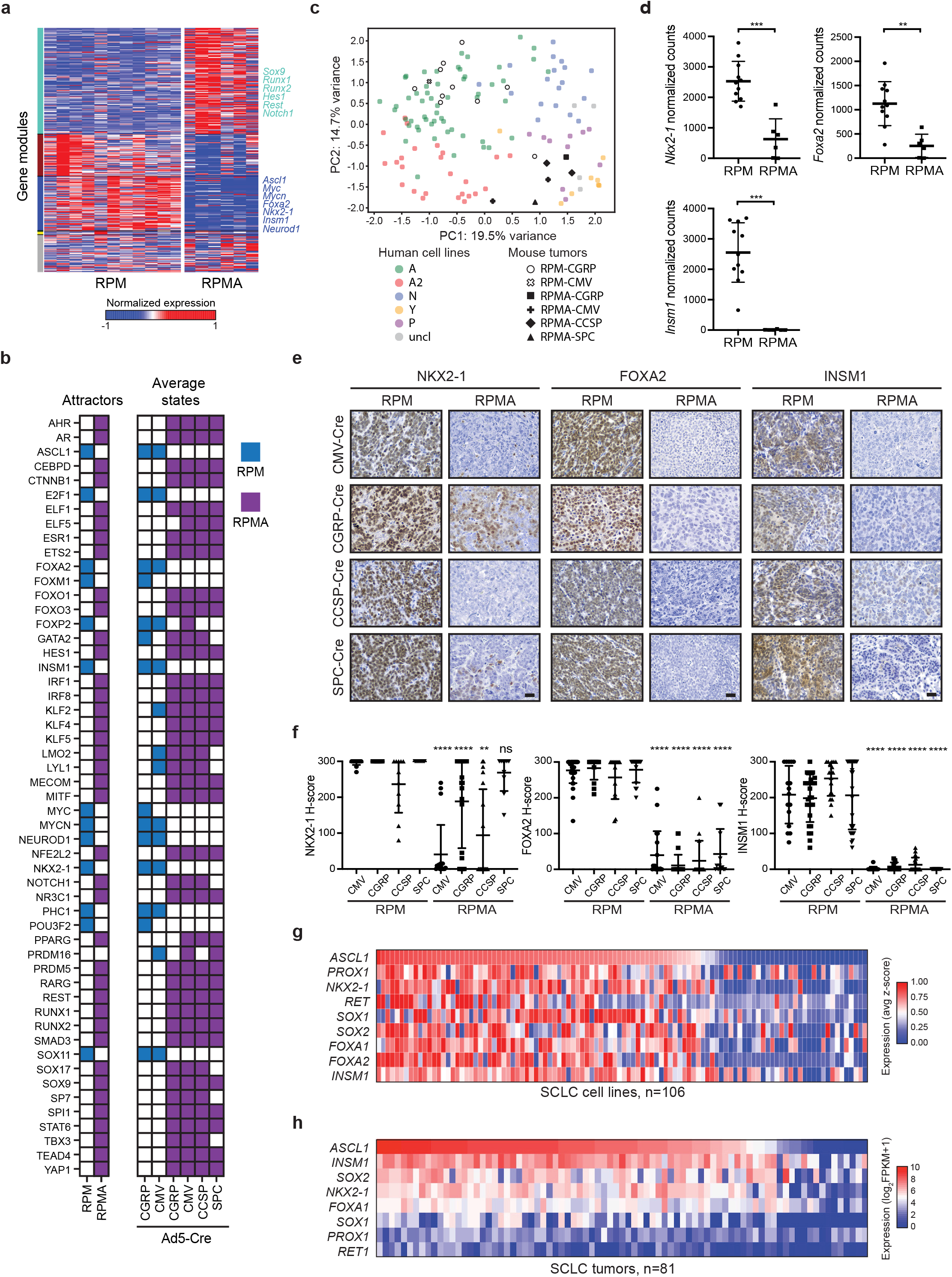
Network analyses predict transcriptional regulators that drive osteosarcoma cell fate upon ASCL1 loss. a) Weighted Gene Coexpression Network Analysis (WGCNA) reveals co-expressed gene modules in RPM and RPMA tumors. b) Binarized average states and found attractors in RPM (blue) and RPMA (purple) tumors initiated with the indicated Cre viruses. For each gene, colored squares are ON and white squares are OFF. c) Principal component analysis comparing bulk RNA-seq expression in human SCLC cell lines to mouse RPM or RPMA tumors initiated with the indicated viruses. Human cell lines were classified into subtypes based on high expression of *Ascl1* (A and A2 variant), *Neurod1* (N), *Pou2f3* (P), or *Yap1* (Y) or were unclassified (uncl). d) Normalized counts of indicated genes from RNA-seq. Data is shown as mean ± SD. Mann-Whitney two-tailed t-test, *** p < 0.001; ** p < 0.01. e) Representative IHC images for indicated antibodies in RPM and RPMA mice infected with cell-type-specific Cre viruses. Scale bar: 25 μm. f) H-Score IHC quantification for indicated proteins in RPM and RPMA mice. Approximately 5-33 tumors were quantified from n = 3-5 mice per condition. Data is shown as mean ± SD. Mann-Whitney two-tailed t-test, **** p < 0.0001; ** p < 0.01; ns = not significant. g) Heat map derived from “SCLC-CellMinerCDB” showing relative expression of SCLC transcription factor genes compared to *ASCL1*. h) Heat map showing relative expression of SCLC transcription factor genes in human tumors from George et al, *Nature*, 2015. See also Supplementary Figure 3.

In our network, we included other known neuroendocrine fate specifiers like INSM1 and transcription factors known to be important in endodermal cell fate and lung adenocarcinoma fate, such as NKX2-1 and FOXA2. While all of these genes were differentially expressed between RPM and RPMA tumors (Fig. 4d), only FOXA2 significantly affected the dynamics of the transcription factor network. IHC analyses confirmed loss of NKX2-1, FOXA2, and INSM1 protein in RPMA compared to RPM tumors, largely independent of initiating cell of origin (Fig. 4e and 4f). Interestingly, NKX2-1 levels remained high particularly in CGRP and SPC-initiated tumors; however, it is noted that the SPC-initiated NKX2-1^+^ tumors appeared to be largely adenocarcinoma. Consistently, *ASCL1* expression is positively correlated with lineage-related transcription factor genes including *NKX2-1, FOXA1, FOXA2*, and *INSM1* in human SCLC cell lines (Fig. 4g) from the SCLC-CellMiner database^23^, with similar results in human tumor data (Fig. 4h). Thus, ASCL1 loss leads to coordinate loss of other key lineage-related transcription factors, which are known to function in the same SCLC super-enhancers^12,24,25^.

### ASCL1 represses non-endodermal cell fates and Notch/Wnt developmental pathways

We next focused on understanding the cell fate and predicted regulators gained upon ASCL1 loss. The lung epithelium is believed to derive largely from endoderm, whereas bone fates are derived from mesoderm or ectoderm^26,27^. Mesenchymal stem cells (MSCs) from mesodermal tissue and neural crest stem (NCS) cells from ectodermal tissue are known to have the potential to become neurons or bone^28,29^. Thus, we examined whether gene signatures for these cell types are enriched in RPM or RPMA tumors. GSEA revealed that both MSC and NC-cell signatures were significantly enriched in RPMA vs RPM tumors (Fig. 5a). Consistent with these results, GSEA revealed that both MSC and NC-cell signatures were significantly enriched in a panel of 109 human SCLC cell lines when comparing *ASCL1*-low versus *ASCL1*-high samples (Fig. 5b). These data suggest that ASCL1 facilitates an endodermal tumor cell fate and represses an MSC/NC-stem-like fate.

**Figure 5:**
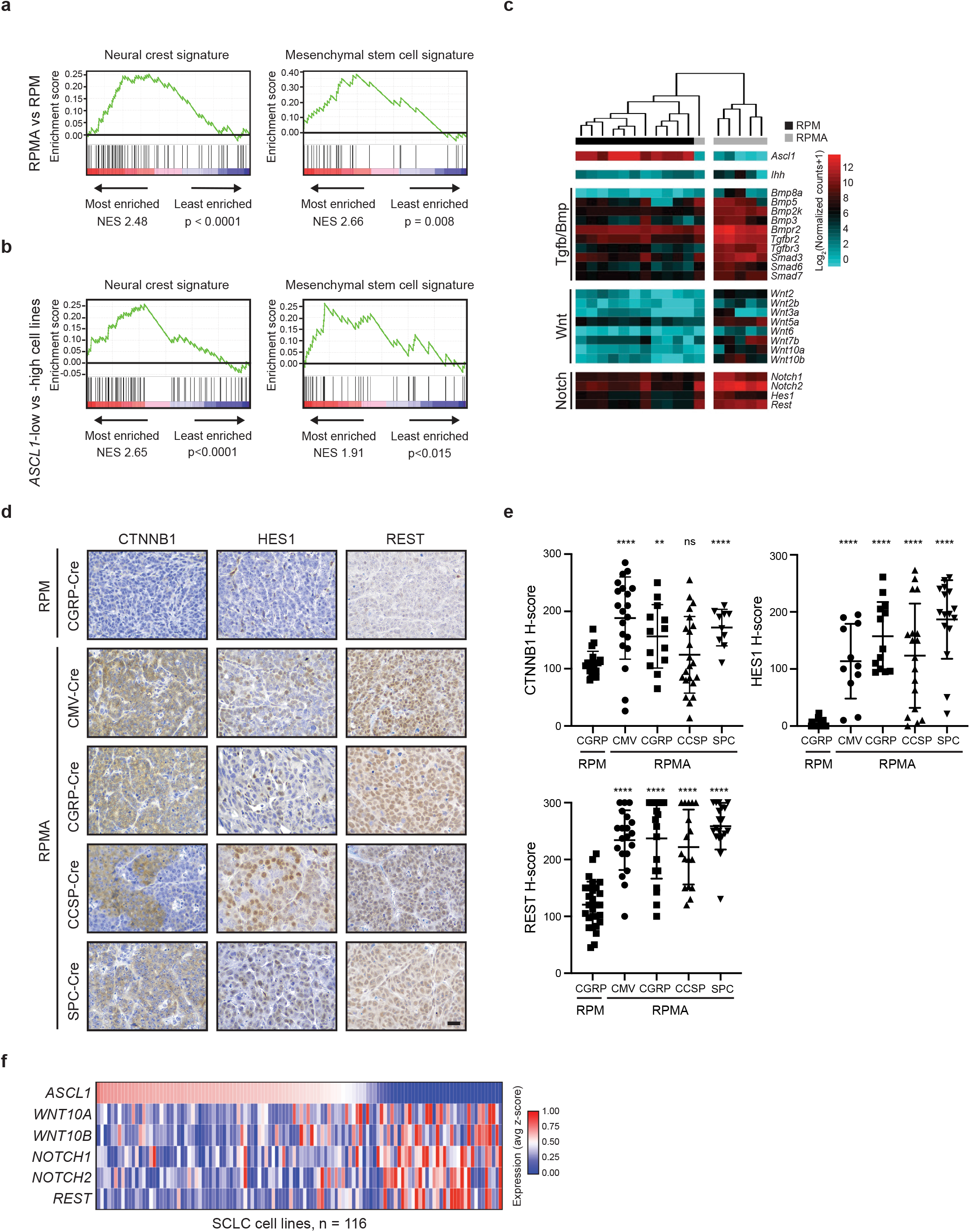
ASCL1 represses non-endodermal cell fates and Notch/Wnt developmental pathways. a) GSEA in RPMA vs RPM tumors compared to neural crest and mesenchymal stem cell signatures. NES and p values indicated in the figure. b) GSEA in *ASCL1*-low vs *ASCL1*-high human cell lines (SCLC-CellMinerCDB) compared to neural crest and mesenchymal stem cell signatures. NES and p values indicated in the figure. c) Heat map showing expression of indicated genes in RPM vs RPMA tumors analyzed by RNA-seq for components of the Indian hedgehog (Ihh), TGFβ, BMP, WNT, and NOTCH pathways. d) Representative IHC for CTNNB1, HES1, and REST in RPM vs RPMA tumors initiated with the indicated cell-type-specific-Cre viruses. Scale bar: 25 μm. e) H-score IHC quantification for indicated antibodies in panel d. Data is shown as mean ± SD. Approximately 11-28 tumors were used for quantification from 3-5 animals per condition. Mann-Whitney two-tailed t-test, **** p < 0.0001; ** p < 0.01; ns = not significant. f) Heat map derived from “SCLC-CellMinerCDB” with expression of selected genes involved in WNT and NOTCH pathways relative to *ASCL1* in human SCLC cell lines.

During development, both MSC and NC-stem-like progenitors have the capacity for bone development given the appropriate developmental signals. Multiple developmental pathways are known to drive bone fate including Indian hedgehog (Ihh), Transforming growth factor-beta (Tgfβ), Bone morphogenic protein (Bmp), Wnt, Notch and others^30^. Numerous components of these pathways were transcriptionally upregulated in RPMA tumors including *Ihh*, Bmp family members (*Bmp3, 5, 2k, 8a, r2*), Tgfβ receptors (*Tgfbr2, 3*), Smads (*Smad3, 6, 7*), Wnt ligands (*Wnt2, 2b, 3a, 5a, 6, 7b, 10a, 10b*), and Notch receptors and target genes (*Notch1, Notch2, Hes1*, and *Rest*) (Fig. 5c). Importantly, network analyses predicted that CTNNB1, HES1, NOTCH1 and REST promote the RPMA fate (Fig. 4b). Thus, we determined Wnt and Notch pathway activity by examining protein levels of targets for which we could identify good antibodies, including CTNNB1 (β-catenin), HES1 and REST (Fig. 5d). Consistent with bioinformatic predictions, CTNNB1, HES1, and REST levels were significantly increased in RPMA compared to RPM tumors regardless of cell of origin (Fig. 5e). Similarly, gene expression data from human SCLC cell lines showed an inverse correlation between *ASCL1* and key Wnt and Notch pathway genes (Fig. 5f). Together, these data suggest that ASCL1 represses an MSC/NC-stem like state as well as Wnt and Notch pathway activity in MYC-driven SCLC.

### ASCL1 represses SOX9 in mouse and human SCLC

In both the developing limb bud mesenchyme and neural crest, SOX9 marks osteoblast progenitors and precedes RUNX2 expression and bone differentiation^30^. RUNX1 also precedes RUNX2 expression in developing bone^31^ and has been suggested to function in chondrogenic lineage commitment of mesenchymal progenitor cells^32^. RUNX1 has been shown to directly regulate multiple genes important for bone development including *Sox9* and *Runx2*^33,34^. RUNX2 and SP7 (i.e. Osterix) are required for differentiation of bone-producing osteoblasts, and RUNX2 normally precedes SP7 expression during bone differentiation^30,35,36^. In previous network analysis, RUNX1, SOX9, and RUNX2 were predicted drivers of the RPMA attractor state (Fig. 4b). Therefore, we sought to determine whether the MSC/NCS-like tumor cells express these factors upon ASCL1 loss. Indeed, while RPM tumors rarely expressed RUNX1, SOX9, or RUNX2, all three factors were dramatically increased in RPMA tumors (Fig. 6a, 6b, and Supplementary Fig. 4a). Interestingly, RUNX1 and SOX9 expression were enriched in non-calcified (soft) RPMA tumors compared to the more differentiated osteosarcoma-like tumors. RUNX2 levels were high in non-calcified (soft) RPMA tumors and remained in more differentiated osteosarcomas. In contrast, SP7 was predominantly expressed in the more differentiated osteosarcoma-like tumors (Fig. 6a, 6b, and Supplementary Fig. 4b). High RUNX2 expression was not restricted to RPMA animals with bone-like lesions by microCT. Rather, we identified multiple RPMA-CCSP and SPC animals with RUNX2^+^ tumors that lacked SP7 expression and did not produce osteoid, suggesting that these tumors represent earlier developmental stages prior to formation of calcified bone. Together, these data are consistent with RUNX1 and SOX9’s established roles preceding RUNX2^+^ and SP7^+^ differentiation during bone development.

**Figure 6:**
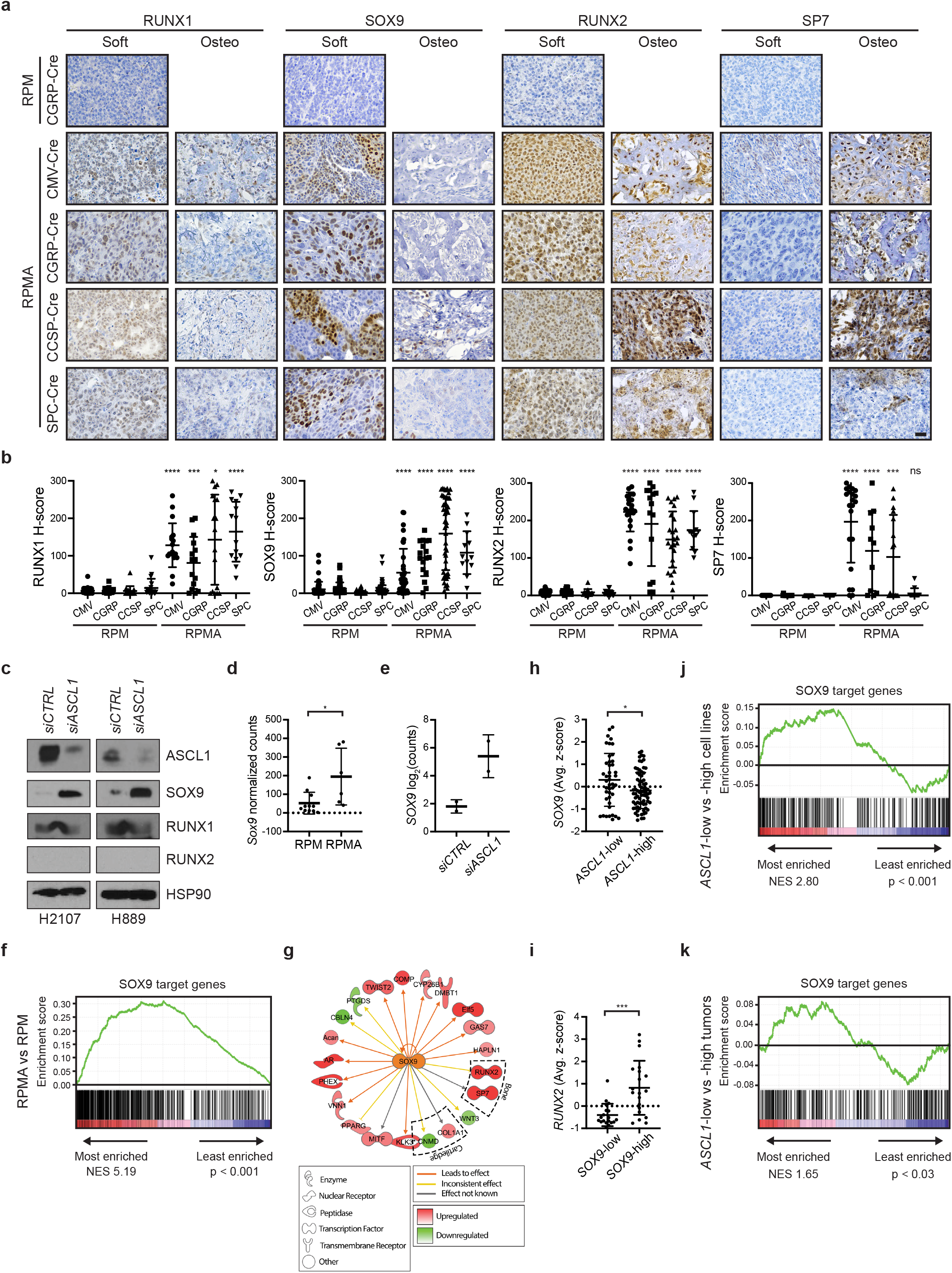
ASCL1 represses SOX9 in mouse and human SCLC tumor cells. a) Representative IHC for RUNX1, SOX9, RUNX2 and SP7 in RPM vs RPMA tumors initiated with the indicated cell-type-specific-Cre viruses. RPMA tumors were classified as non-calcified tumors (soft) or osteosarcomas (osteo). Scale bar: 25 μm. b) H-score quantification for the indicated proteins in RPM vs RPMA tumors from panel a. Data is shown as mean ± SD. Approximately 11-85 tumors from 3-6 mice per condition were quantified. Mann-Whitney two-tailed t-test, **** p < 0.0001; *** p < 0.001; * p < 0.05; ns = not significant. c) Representative immunoblot following 72 hr treatment with control (*CTRL*) or *ASCL1* siRNAs in the indicated human SCLC cell lines. HSP90 serves as loading control. d) *Sox9* expression as normalized counts by RNA-seq from lung tumors in indicated GEMMs. Mean ± SD. Two-tailed t-test, * p < 0.05. e) Expression of *SOX9* by RNA-seq from human SCLC cell line NCI-H2107 treated for 72 hr with *ASCL1* or control *(CTRL)* siRNAs performed in biological duplicates. Results are reported as Log2 normalized counts with mean ± SD. f) GSEA for expression of SOX9 target genes from Larsimont et al, *Cell Stem Cell*, 2015, in RPMA vs RPM tumors. NES and p values indicated in the figure. g) Predicted SOX9 target genes enriched or depleted in RPMA vs RPM tumors by IPA. h) Expression of *SOX9* in human SCLC cell lines grouped by *ASCL1* expression levels in “SCLC-CellMinerCDB.” Data is shown as average z-score ± SD. Mann-Whitney two-tailed t-test, * p < 0.05. i) Expression of *RUNX2* in human SCLC cell lines grouped by *SOX9* expression levels in “SCLC-CellMinerCDB.” Data is shown as average z-score ± SD. Mann-Whitney two-tailed t-test, *** p = 0.005. j) GSEA for expression of SOX9 target genes from Larsimont et al, *Cell Stem Cell*, 2015, in *ASCL1-low* vs *ASCL1*-high human SCLC cell lines from “SCLC-CellMinerCDB.” NES and p values indicated in the figure. k) GSEA for expression of SOX9 target genes from Larsimont et al, *Cell Stem Cell*, 2015, in *ASCL1*-low vs *ASCL1*-high human tumors from George et al, *Nature*, 2015. NES and p values indicated in the figure. See also Supplementary Figure 4.

To determine whether ASCL1 repression of SOX9, RUNX1, and RUNX2 is conserved in human SCLC cells, we knocked down *ASCL1* in human SCLC cell lines (NCI-H2107 and NCI-H889) using siRNAs and examined SOX9, RUNX1 and RUNX2 expression by immunoblot. SOX9 was induced upon *ASCL1* knockdown in both cell lines, while RUNX1 levels did not change, and we did not detect induction of RUNX2 (Fig. 6c). *Sox9/SOX9* is induced transcriptionally in RPMA tumors and human SCLC cell lines, respectively, suggesting that ASCL1 represses its expression (Fig. 6d and 6e). Analysis of ASCL1 ChIP-seq data did not identify significant ASCL1 binding sites near *SOX9* (Supplementary Fig. 4c and 4d), suggesting that repression of *SOX9* by ASCL1 is likely indirect. GSEA showed that established SOX9 target genes were significantly upregulated in RPMA compared to RPM tumors (Fig. 6f), suggesting that SOX9 activity is increased in RPMA tumors. Furthermore, Ingenuity Pathway Analysis (IPA) identified SOX9 as a top transcriptional regulator implicated in the transcriptional differences between RPMA and RPM tumors (Fig. 6g and Supplementary Table 2). IPA also predicted other key transcription factor regulators that distinguish RPMA from RPM such as RUNX2, REST, NKX2-1, FOXA2, NOTCH1, and CTNNB1 (Supplementary Table 2), which we verified herein. In human SCLC cell lines, *SOX9* expression was enriched in samples that were *ASCL1*-low (Fig. 6h), and high *SOX9* expression also correlated with increased *RUNX2* expression (Fig. 6i). GSEA revealed that known SOX9 target genes were significantly enriched in *ASCL1*-low human cell lines (Fig. 6j), and in *ASCL1*-low human SCLC tumors (Fig. 6k). Together these data reveal that ASCL1 represses a SOX9^+^ MSC or NCS-like state that has the capacity for bone differentiation.

## Discussion

The PNEC has been accepted as a major cell of origin for SCLC. This is consistent with multiple similarities between SCLC and PNECs, and with the capacity of PNECs for SCLC transformation in multiple mouse models^18,19,37^. Our data here suggest that specific genetic alterations, namely MYC expression in the context of *Rb1* and *Trp53* loss, can promote SCLC in other cells of origin, including club and alveolar cells, and potentially other cell types as well. It is possible that *Ccsp* and *Spc* are targeting CCSP/SPC-double-positive bronchioalveolar stem cells (BASCs)^38^. While club cells in RP and RPR2 mice were relatively refractory to SCLC development^18,19^, targeting club cells in RPM mice led to SCLC with short latency, comparable to tumor initiation in NE cells. This suggests that club cells may be particularly susceptible to MYC-mediated SCLC transformation. Recent studies suggest that tuft and basal cells may also serve as additional cells of origin in SCLC^8,20^. Together with recent findings^15^, these data are consistent with the notion that cell of origin, genetics, and tumor cell plasticity can determine SCLC phenotype. Moreover, SCLC diagnosis in the clinic does not necessarily mean that tumors arose in NE cells, and this may need to be considered as we develop predictive models for tumor evolution.

Regardless of the cell of origin, RPM tumors demonstrate NE features at the earliest stages of development, suggesting that NE tumors can arise from non-neuroendocrine cells. This suggests that multiple differentiated cells in the adult lung have the capacity to de-differentiate into a neuroendocrine fate. It is notable, however, that some differentiated cell types like AT2 cells appear relatively refractory to transformation even in the context of profound oncogenic changes like *Rb1/Trp53/Myc;* in contrast, AT2 cells are remarkably sensitive to transformation by MAPK pathway activation, as observed upon expression of mutant EGFR or KRAS in GEMMs. Tumor cell fate plasticity can also be appreciated in the clinic as it is known that EGFR-driven lung adenocarcinomas (believed to arise in AT2 cells) can convert to SCLC upon resistance to EGFR inhibitors^39,40^. Approximately 20-30% of prostate adenocarcinoma that acquires resistance to androgen therapy also converts to a neuroendocrine fate through transdifferentiation^41,42^, and these tumors frequently harbor *RB1* and *TP53* loss. Finally, alteration of RB1, TP53, MYC, BCL2, and AKT in prostate or lung basal epithelial cells in vitro can lead to neuroendocrine tumors in xenografts, resembling neuroendocrine prostate cancer and SCLC^20^. These observations illustrate the potent capacity of oncogenic changes to alter cell fate, and in particular, how RB1, TP53 and MYC cooperate to promote NE fate.

Given ASCL1’s role as a lineage-specific oncogene in SCLC that is highly expressed in a significant fraction of tumors, this has prompted its consideration as a therapeutic target^43^. This notion has been supported by studies showing that genetic deletion of *Ascl1* in classic GEMMs abolishes tumor formation^12^. However, here we find that ASCL1 is not required for MYC-driven tumor development in the RPM model, even though it does appear to be required for neuroendocrine cell fate. An important caveat in both studies is that *Ascl1* is deleted at the time of tumor initiation. It remains to be tested whether ASCL1 is necessary for the growth of established tumors, with and without MYC expression, which will require the development of more advanced conditional GEMM systems. ASCL1 is lowly expressed in MYC-driven human SCLC^6,10^, together suggesting that ASCL1 inhibition may not be sufficient to block the growth of MYC-driven SCLC. Our data suggest that loss of ASCL1 may simply convert SCLC to an alternative cell fate. Indeed, chemotherapy-relapsed SCLC was found to exhibit significantly reduced ASCL1 expression^44^, suggesting ASCL1 may not be required for SCLC progression. Interestingly, chemotherapy-relapsed tumors had evidence for a gain in WNT pathway activity^44^, consistent with our observations here that imply an antagonistic relationship between ASCL1 and WNT signaling in SCLC.

We were surprised to find that genetic disruption of *Ascl1* led to conversion of SCLC to an osteosarcoma-like fate. Both *RB1* and *TP53* alterations are remarkably common in osteosarcomas (*RB1*, 70-90%; *TP53*, 50-70%)^45,46^. In addition, loss of *Rb1* and *Trp53* in osteoblasts or MSCs in the mouse promotes tumors that highly resemble human osteosarcoma, including high expression of RUNX2^45,47,48^. During osteogenesis, RUNX2 has been shown to bind pRB1 and HES1^49–51^, and we observe upregulation of HES1 and other indicators of active NOTCH signaling in RPMA tissue. ASCL1 and NOTCH/REST exhibit a mutually antagonistic relationship in multiple contexts^52,53^. In SCLC, ASCL1 and Notch have an established relationship whereby Notch signaling promotes non-neuroendocrine fate at least partially through induction of the transcriptional co-repressor REST^54^. We observe induction of NOTCH/REST in RPMA tumors, and we speculate that this event is key to inhibiting NE fate. As NOTCH, WNT and other developmental pathways impinge upon RUNX2 to drive bone fate, future studies will be required to elucidate the signals that promote bone differentiation in the context of ASCL1 loss.

Our findings demonstrate that alterations in *Rb1*, *Trp53* and *Myc* cooperate to dedifferentiate tumor cells to a state that has the potential to be neuroendocrine in the presence of ASCL1, and bone-like in its absence. One outstanding question prompted by these findings regards the nature of this dedifferentiated cell state; during development, the lung epithelium is believed to arise from endodermal progenitors, whereas bone is derived from mesoderm or ectoderm. Therefore, is there a developmental cell type with NE and bone potential that could be related to SCLC? At least two cell types are known to have the capacity for neural and bone fates during development, MSCs (whose origin is not well understood) and neural crest (NC) stem cells, which can give rise to both neurons and facial bone^30,55,56^. Gene expression analyses of RPMA tumors suggest that they harbor MSC and NC stem cell-like gene expression signatures. Other “small, round blue cell” tumors such as Ewing’s sarcoma have also been shown to resemble MSCs^57^. While this issue requires further study, it suggests the surprising notion that in the absence of ASCL1, SCLC can dedifferentiate to a SOX9^+^ stem-like cell that precedes commitment to endodermal lineages.

These findings have led us to question whether SCLC can evolve to a bone-like state in patients. A slow-growing bone phenotype would be under *strong negative selective pressure* compared to rapidly-growing SCLC, and therefore likely to be an extremely rare event. However, there are rare case reports of patients developing carcinoids with ossification^58,59^ and extraskeletal osteosarcoma (ESOS) in the lung^60^, including one ESOS following chemotherapy treatment of SCLC^61^. It is tempting to speculate that treatments that block ASCL1 potently in the clinic, could push tumor evolution to a bone-like fate, or more likely, to a faster-growing dedifferentiated MSC or NC stem cell-like state. Further study of chemotherapy-relapsed tissue will be required to address this possibility.

Finally, we observed that *Neurod1* was one of the most downregulated genes upon ASCL1 loss in MYC-driven SCLC. This suggests the intriguing possibility that the NEUROD1^+^ subtype of SCLC requires ASCL1 for development. It has been debated whether the NEUROD1^+^ SCLC subtype arises in the lung^12^; both our recent data^15^ and that shown here suggest that the NEUROD1^+^ subtype is promoted by MYC but temporally follows ASCL1 expression. There are developmental contexts where ASCL1 expression precedes NEUROD1 such as in the olfactory epithelium and during adult hippocampal neurogenesis^62,63^. Thus, we speculate that NEUROD1 expression follows ASCL1 during MYC-driven tumor progression in SCLC. Together, our study suggests that cancer cells demonstrate remarkable plasticity and capacity to evolve to early developmental programs.

## Methods

### Mice

Animal experiments were approved by the Institutional Animal Care and Use Committee at the University of Utah and performed in compliance with all relevant ethical regulations. *Rb1^fl/fl^;Trp53^fl/fl^;MycT58A^LSL/LSL^* (RPM, JAX #029971) mice were previously described^6^. *Rb1^fl/fl^;Trp53^fl/fl^; Rbl2^/fl^;Ascl1^fl/fl^* mice were provided by J.E. Johnson^12^. Sperm was collected from mice carrying the conditional *Ascl1* allele^64^ and used to cross to our RPM strain through in vitro fertilization. Anesthetized mice at 6-8 weeks of age were infected by intratracheal instillation^65^ with 1×10^8^ plaque-forming units of adenovirus carrying Cre recombinase (University of Iowa). Viruses include: Ad-CMV-Cre (VVC-U of Iowa-5), Ad-CGRP-Cre (VVC-Berns-1160), Ad-CCSP/CC10-Cre (VVC-Berns-1166), and Ad-SPC-Cre (VVC-Berns-1168). Viral infections were performed in a Biosafety Level 2+ room following guidelines from the University of Utah Institutional Biosafety Committee. Male and female mice were divided equally for all experiments. For survival studies, mice were sacrificed due to tumor burden or visibly labored breathing. Mice were euthanized by CO2 asphyxiation followed by necropsy.

### MicroCT imaging

Mice were anesthetized with isoflurane and imaged using a small animal Quantum FX or Quantum GX2 microCT (Perkin Elmer). Quantum FX images were acquired with 34 sec scans at 90 kV, 160 μA current, and reconstructed at a 45 μm voxel size. Quantum GX2 images were acquired with 18 sec scans at 90 kV, 88 μA current, and reconstructed at a 90 μm voxel size.

For improved imaging of bone, selected animals were imaged for 2 min on the Quantum GX2 at 90 kV, 88 μA current, and reconstructed at a 90 μm voxel size. Frequency of imaging ranged from twice/week to once/month depending on the tumor growth rate. Resulting images were processed with Analyze 11.0 or 12.0 software (Analyze Direct) as previously described^6^. For bone analysis, acquired images were viewed with the Quantum GX2 Simple Viewer.

### Immunohistochemistry

At necropsy, lungs were inflated with PBS, fixed overnight in 10% neutral buffered formalin, and then transferred to 70% ethanol. Samples requiring demineralization were incubated in Rapid-Cal Immuno Decal Solution (6089, BBC Chemical) prior to paraffin embedding. Tissue was cut into 4 μm sections and stained with H&E to assess tumor pathology, or with specific antibodies. Slides were deparaffinized in Citrisolv clearing agent (22-143-975, Fisher Scientific), and rehydrated in an ethanol dilution series. Heat-mediated antigen retrieval was performed in a pressure cooker for 15 min using either 0.01 M citrate buffer pH 6.0 or 0.01 M TE buffer pH 9.0, as suggested by the antibody manufacturer. Endogenous peroxidases were blocked by 10 min incubation with 3% H_2_O_2_, prior to 30 min blocking in 5% goat serum/PBST (PBS containing 0.1% Tween-20). Primary antibodies were diluted in either 5% goat serum/PBS-T or SignalStain Boost IHC Detection Reagent (8114S, Cell Signaling Technology (CST)), as recommended by the manufacturer. Slides were incubated in primary antibody overnight at 4°C using the Sequenza coverslip staining system (ThermoFisher Scientific). The next day, slides were incubated for 30 min at room temperature with VectaStain secondary antibody (PK-4001, BMK-2202, Vector Laboratories), followed by 30 min incubation in ABC reagent and development with DAB (SK-4100, Vector Laboratories). Slides were counterstained with hematoxylin and mounted with a Shandon ClearVue coverslipper (Thermo Scientific). Primary antibodies include the following: ASCL1 (556604, BD Pharmingen, 1:200), CCSP (AB07623, Millipore Sigma, 1:2000), CGRP (C8198, Sigma-Aldrich, 1:250), CTNNB1 (9587S, CST, 1:250), FOXA2 (ab108422, Abcam, 1:1000), HES1 (11988S, CST, 1:600), MYC (sc-764, Santa Cruz, 1:150), NEUROD1 (ab109224, Abcam, 1:150), NKX2-1 (ab76013, Abcam, 1:250), POU2F3 (HPA019652-100UL, Sigma-Aldrich, 1:300), REST (ab202962, Abcam, 1:100), RUNX1 (25315-1-AP, Proteintech, 1:250), RUNX2 (ab192256, Abcam, 1:1000), SOX9 (ab185966, Abcam, 1:1000), SP7 (ab209484, Abcam, 1:1000), SPC (AB3786, Millipore Sigma, 1:2000), SYP (RB1461P1, Thermo Fisher Scientific, 1:200), UCHL1 (HPA005993-100UL, Sigma-Aldrich, 1:250), and YAP1 (14074S, CST, 1:400). Images were acquired on a Nikon Ci-L LED Microscope with DS-Fi3 Camera. Selected slides were digitally scanned with a Pannoramic Midi II (3DHISTECH) slidescanner. Staining was manually quantified by H-score on a scale of 0-300 taking into consideration percent positive cells and staining intensity as described^66^, where H Score = % of positive cells multiplied by intensity score of 0-3. For example, a tumor with 80% positive cells with high intensity of 3 = 240 H-Score. For SOX9 expression, scanned IHC slides were quantified by H-score using CaseViewer software (3DHISTECH) and an automated nuclear quantification program after manual identification of tumor regions.

### Mouse tumor RNA-seq

RNA isolation from ~15 mg flash-frozen RPM (n=11), un-calcified RPMA (n=7), and RPR2 (n=6) primary tumors was performed using RNeasy Mini Kit (Qiagen) with the standard protocol. RNA from RPM (n=11) and RPR2 (n=2) tumors was subject to library construction with the Illumina TruSeq Stranded mRNA Sample Preparation Kit (RS-122-2101, RS-122-2102) according to the manufacturer’s protocol. RNA from RPR2 (n=4) and RPMA (n=7) tumors was subject to library construction with the Illumina TruSeq Stranded Total RNA Library Ribo-Zero Gold Prep kit (RS122-2301) according to the manufacturer’s protocol. Chemically denatured sequencing libraries (25 pM) from RPM (n=11) and RPR2 (n=6) tumors were applied to an Illumina HiSeq v4 single read flow cell using an Illumina cBot. Hybridized molecules were clonally amplified and annealed to sequencing primers with reagents from an Illumina HiSeq SR Cluster Kit v4-cBot (GD-401-4001). Following transfer of the flowcell to an Illumina HiSeq 2500 instrument (HCSv2.2.38 and RTA v1.18.61), a 50-cycle single-end sequence run was performed using HiSeq SBS Kit v4 sequencing reagents (FC-401-4002). Chemically denatured sequencing libraries (1.15 nM) from RPMA tumors (n=7) were applied to an Illumina NovaSeq S2 flow cell using the standard workflow. Hybridized molecules were clonally amplified and annealed to sequencing primers with reagents from an Illumina NovaSeq 6000 S2 Reagent Kit (20012861). Following transfer of the flowcell to an Illumina NovaSeq 6000 instrument (20012850), a 50-cycle paired-end sequence run was performed.

### Bioinformatics

Fastq raw count files were aligned in the R statistical environment (v3.6). The mouse GRCm38 FASTA and GTF files were downloaded from Ensembl release 94 and the reference database was created using STAR version 2.6.1b^67^ with splice junctions optimized for 50 base pair reads. Optical duplicates were removed using clumpify v38.34 and adapters were trimmed using cutadapt 1.16^68^. The trimmed reads were aligned to the reference database using STAR in two pass mode to output a BAM file sorted by coordinates. Mapped reads were assigned to annotated genes in the GTF file using featureCounts version 1.6.3^69^. The output files from cutadapt, FastQC, Picard CollectRnaSeqMetrics, STAR and featureCounts were summarized using MultiQC^70^ to check for any sample outliers. To remove sources of unwanted variation from tumor RNA-seq library preparation and sequencing platforms, all non-coding features, histones, and ribosomal RNAs were removed for downstream analyses. The featureCount output files were combined into a single raw count matrix. Differentially expressed genes (DEGs) were identified using a 5% false discovery rate with DESeq2 version 1.24.0^71^ (Supplementary Table 1). These data have been deposited in NCI GEO: GSE155692. PCA was performed on the first two principle components using the regularized log count (rlog) values of the top 500 variable genes. Log2(+1)-transformed, normalized intensity values were obtained to create heat maps with unsupervised hierarchical clustering. A previously-described NE marker gene set^6^ was used to compare RPMA (n=6) and RPM (n=11) tumor expression. Heat maps were generated using Pheatmap version 1.0.12. Semantic similarity-based scatterplot visualizations of enriched and depleted GO Biological Processes in RPMA vs RPM tumors were created with the REViGO tool using GO Biological Processes input lists and corresponding p-values^72^. Scatterplot view visualizes the GO terms in a “semantic space” where the more similar terms are positioned closer together. The color of the bubble and size reflects the log 10 p-value obtained in the Panther analysis. Input GO Biological Processes and log10 p-values were generated using Panther v14 GO Enrichment Analysis tool for biological processes^73^ on enriched (>2.5 log2FC by DESeq2) or depleted (<-2.5 log2FC by DESeq2) genes in RPMA vs RPM tumors. REViGO and Panther GO Enrichment Analysis details and curated input gene lists are presented in Supplementary Table 1.

### GSEA

Gene set enrichment analysis (GSEA) was performed using GSEA version 3.0 software with default parameters, classic enrichment, inclusion gene set size between 15 and 1000, and the phenotype permutation at 1,000 times. GSEA was performed on a pre-ranked gene list representing log2 fold change of RPMA vs RPM tumor bulk RNA-seq expression values (Supplementary Table 1). Normalized enrichment scores and p values are shown below each respective GSEA plot in the figures. A catalog of functional gene sets from Molecular Signature Database (MSigDB, version 6.2, July 2018, www.broad.mit.edu/gsea/msigdb/msigdb_index.html) was used for the ‘‘GO Bone Development”, “Mesenchymal stem cell” and “Neural crest signature” gene sets. Gene sets for “ASCL1 Targets by ChIP-Seq” and “NEUROD1 Targets by ChIP-Seq” represent conserved transcriptional targets identified by ChIP-Seq from mouse SCLC tumor and human cell line SCLC models. The ASCL1 conserved target gene list was previously published (GEO: GSE69398)^12^. The NEUROD1 conserved target gene list was derived from published ChIP-seq data from human SCLC cell lines (GEO: GSE69398)^12^ and NEUROD1 ChIP-seq data from RPM mouse tumors described in this study. The “SOX9 target genes” list was derived from published mouse basal cell carcinoma ChIP-seq data (GEO: GSE68755)^74^. Details of the curated gene sets are presented in Supplementary Table 1.

### ASCL1 and NEUROD1 ChIP-seq

Preparation of mouse lung tumors for ChIP-seq was performed as previously described^7^. Briefly, 20 million cells or ~250 μg of mouse tumor per ChIP were crosslinked in 1% formaldehyde for 10 min at room temperature. Crosslinking was stopped with 125 mM glycine and nuclei were extracted. Chromatin was sonicated using an Epishear Probe Sonicator (Active Motif) for 4 min at 40% power. ASCL1 antibody (556604, BD Pharmingen), H3K27Ac antibody (39133, Active Motif), or NEUROD1 antibody (sc-1084, Santa Cruz Biotechnology) were used for IP. An input sample of each RPM tumor served as the control. Libraries were sequenced on an Illumina HiSeq 2500 as single-end 50 bp reads to a minimum depth of 35 million reads per sample. Reads were aligned to the mm10 build of the mouse genome with bowtie^75^ using the following parameters: -m 1 -t --best -q -S -l 32 -e 80 -n 2. Peaks were called with MACS2^76^ using a p value cutoff of 10^-10^ and the mfold parameter bounded between 15 and 100. For visualization, MACS2 produced bedgraphs with the –B and –SPMR options. Binding profiles were visualized using Integrated Genome Viewer (IGV version 2.6.3) aligned to mm10 genome build. ASCL1 and NEUROD1 mouse ChIP-seq data are available on GEO: GSE155692. H3k27Ac mouse ChIP-seq data are available on GEO: GSE142496^7^.

Human ChIP-seq for ASCL1 was performed as described in Borromeo et al., 2016 and data are published in GEO with accession number GEO: GSE69398. Binding profiles of human ASCL1 ChIP-seq were visualized using Integrated Genome Viewer (IGV version 2.6.3) aligned to hg19 human genome build.

### Weighted Gene Coexpression Network Analysis

WGCNA^77^ was performed on RPM (n=11) and RPMA (n = 6) RNA seq data using the R package ‘WGCNA’. Low variance genes, with coefficient of variation < 0.2 and median absolute deviation < 1, were removed, resulting in 15,286 genes across 17 samples. A signed network of genes, such that only positively correlated genes will be grouped into gene modules (negative correlations are given a score of 0), was constructed as follows. WGCNA’s pickSoftThreshold function was used to determine the exponent of the correlation coefficients matrix that best produces a scale free network when the matrix is used as weights of network connections. A threshold of 15 was chosen to produce this weights matrix, with the correlation function “Pearson.” A topological overlap matrix was then generated based on the overlap in connections for two genes. This gave a measure of distance between each pair of genes, which was used in average-linkage hierarchical clustering. The ‘cutTreeDynamic’ function was used, with minimum cluster size of 100 to generate clusters, or modules, of coexpressed genes. WGCNA’s ‘sampledBlockwiseModules’ was then used to increase robustness of gene modules, which merges or removes genes from modules that are not distinct enough in a majority of the resampled module labels. Differentially-expressed gene modules were determined using an ANOVA statistical test between RPM and RPMA tumors. Three gene modules were significant.

### Network Structure Inference

Network structure was inferred as described in Wooten et al (2019)^22^. Briefly, transcription factors were filtered to include those differentially expressed between RPM and RPMA tumors with Log Fold Change > 1.5 and adjusted p value < 0.05. Genes were then filtered further to include only those most central to each differentially-expressed gene module, calculated as a module membership value (kME) from the WGCNA analysis. Lastly, expert knowledge was used to add any transcription factors that were deemed central to RPM/RPMA cell identity, including FOXA2, SOX9, and CTNNB1. Network connections were added using the Enrichr tool from the Ma’ayan laboratory^78,79^, which mines various ChIP-seq databases for TF-gene interactions including CHEA, ENCODE, and TRANSFAC. Gene nodes with no downstream nodes (child nodes) were removed, as they do not affect the dynamics of the network, except NKX2-1, INSM1, and SP7, which were kept as biomarkers.

### Network Rule Inference and Attractors

Network rules of interaction were calculated using BooleaBayes as previously described^22^, using the 11 RPM and 6 RPMA samples. For each TF node in the network, a rule of expression was determined for any ON/OFF combination of parent nodes. BooleaBayes simulations consisted of asynchronous directed random walks, with probabilities of transition between states determined by TF rules. By simulating the network starting at each sampled state, we found pseudo-attractors (here, referred to as “attractors) as previously defined^22^. Searches for attractor states from each initial condition were cut-off at a TF-basin radius of 4 steps, and the phenotypes of any attractors were determined by distance to initial condition. Two attractors were found using this method, one for each phenotype. Driving transcription factors for each attractor, or those whose silencing have a destabilizing effect on the attractor, were found using *in silico* perturbations^22^.

### ASCL1 knockdown and immunoblot

Human classic SCLC cell lines NCI-H2107 and NCI-H889 were obtained from ATCC. Cells were transfected with human *ASCL1* stealth siRNA (129901-HSS100744, Thermo-Fisher) using X-tremeGENE siRNA Transfection Reagent (4476115001, Sigma) according to the manufacturer’s protocol. Cells were harvested 72 hr after transfection for analysis by immunoblot. Cell pellets were flash frozen and stored at −80°C until use. Total protein lysates were prepared as previously described, separated via SDS-PAGE and transferred to a PVDF membrane^80^. Membranes were blocked for 1 hr in 5% milk followed by overnight incubation with primary antibodies at 4°C. Membranes were washed for 4 x 10 min at RT in TBS-T. Mouse and rabbit HRP-conjugated secondary antibodies (711-035-152 and 115-035-205, Jackson ImmunoResearch, 1:10,000) were incubated for 1 hr in 5% milk at RT followed by washing 4 x 10 min at RT in TBS-T. Membranes were exposed to WesternBright HRP Quantum substrate (Advansta) and detected on Hyblot CL film (Denville Scientific Inc). Primary antibodies include: ASCL1 (556604, BD Pharmingen, 1:300), SOX9 (ab185966, Abcam, 1:1000), RUNX1 (25315-1-AP, Proteintech, 1:1000), RUNX2 (12556, CST, 1:1000), with HSP90 (4877, CST, 1:5000) as a loading control.

### PCR for Cre recombination efficiency

Approximately 15 mg of flash-frozen tumor tissue was cut on dry ice and processed with the Qiagen DNeasy kit to isolate tumor genomic DNA. DNA concentrations were measured on a BioTek Synergy HT plate reader. Equal quantities of tumor genomic DNA were amplified by PCR with GoTaq (M7123, Promega) using the gene-specific primers listed. For *Ascl1*: Common Fwd (MF1) 5’-CTACTGTCCAAACGCAAAGTGG-3’; WT Rev (MR1) 5’-GCTCCCACAATCCTCGTAAAGA-3’; Flox Rev (VR2) 5’-TAGACGTTGTGGCTGTTGTAGT-3’; and Recombined Fwd (5’ UTR Fwd) 5’-AACTTTCCTCCGGGGCTCGTTTC-3’. PCR conditions were: 94 deg 5 min, 30 cycles of (94 deg 1 min, 64 deg 1.5 min, 72 deg 1 min), 72 deg 10 min, hold at 4 deg. Expected band sizes are as follows: 5’ UTR Fwd-VR2 (recombined allele) = ~700 - 850 bp, MF1-VR2 (floxed allele) = 857 bp, MF1-MR2 (wild-type allele) = 411 bp.

Primers to detect *Rb1* recombination include the following: D1 5’-GCAGGAGGCAAAAATCCACATAAC-3’, 1lox 5’ 5’-CTCTAGATCCTCTCATTCTTCCC-3’, and 3’ lox 5’-CCTTGACCATAGCCCAGCAC-3’. PCR conditions used were 94 deg 3 min, 30 cycles of (94 deg 30 sec, 55 deg 1 min, 72 deg 1.5 min), 72 deg 5 min, hold at 4 deg. Expected band sized were ~500 bp for the recombined allele, and 310 bp for the floxed allele.

Primers to detect *Trp53* recombination include the following: A 5’-CACAAAAACAGGTTAAACCCAG-3’, B 5’-AGCACATAGGAGGCAGAGAC-3’, and D 5’-GAAGACAGAAAAGGGGAGGG-3’. PCR conditions were 94 deg 2 min, (30 cycles of 94 deg 30 sec, 58 deg 30 sec, 72 deg 50 sec), 72 deg 5 min, hold at 4 deg. Expected band sizes were 612 bp for the recombined allele, and 370 bp for the floxed allele. PCR products were run on 1-2% agarose/TAE gels containing ethidium bromide and images were acquired using an Azure Biosystem C200 imager.

### Statistics

Statistical analysis was performed with GraphPad Prism 8. Mouse survival studies were analyzed with Mantel-Cox log-rank tests. Immunohistochemistry quantification and normalized RNAseq counts were compared by non-parametric two-tailed Mann-Whitney t-tests. Measurements were taken from distinct tumors in multiple mice for each experiment, as indicated in the figure legend. P values < 0.05 were considered statistically significant. Error bars represent mean ± standard deviation unless otherwise specified.

## Acknowledgements

We appreciate Dr. Anton Berns for permission to use Ad-Cre viruses deposited at University of Iowa. We are grateful to the late Dr. Adi Gazdar, who provided insightful pathological observations for this study. We thank R. Dahlgren and L. Houston for administrative assistance and members of the Oliver lab, especially D. Hansen, P. Ballieu, and A. Micinski for mouse colony management. We acknowledge support from the National Cancer Institute (NCI) of the National Institutes of Health (NIH) under award P30CA042014 to Huntsman Cancer Institute for the use of core facilities and shared resources, including Biorepository and Molecular Pathology, and High-Throughput Genomics and Bioinformatic Analysis. This work was supported by the NIH NCI (R21-CA216504-01A1; U01-CA231844; U01-CA213338).

## Author Contributions

Conceptualization: R.R. Olsen and T.G. Oliver

Data Acquisition: R.R. Olsen, D.W. Kastner, A.S. Ireland, K. Pozo, S. Wait, M.R. Guthrie, C.P. Whitney, D. Soltero

Data Analysis and Interpretation: R.R. Olsen, D.W. Kastner, A.S. Ireland, S.M. Groves, B.L. Witt, V. Quaranta, T.G. Oliver

Writing – Original Draft: R.R. Olsen, A.S. Ireland, T.G. Oliver

Writing – Review and Editing: R.R. Olsen, A.S. Ireland, S.M. Groves, V. Quaranta, J.E. Johnson, T.G. Oliver

Funding Acquisition: T.G. Oliver

Resources: J.E. Johnson

Supervision: V. Quaranta, J.E. Johnson, T.G. Oliver

## Resource availability

## Corresponding author

Further information and requests for resources and reagents should be directed to and will be fulfilled by the corresponding author, Trudy G. Oliver.

## Materials availability

The RPM mice used in this study are deposited at The Jackson Laboratory, JAX #029971. RPMA mice are available upon request with an MTA.

## Data availability

All software is commercially available or cited in previous publications. Mouse RNA-seq and ChIP-seq data unique to this study are deposited in NCBI GEO: GSE155692. Associated raw data is available for Fig. 3–6 and is provided in Supplementary Tables 1-2.

