## Supplementary Figures for "ASCL1 represses a latent osteogenic program in small cell lung cancer in multiple cells of origin"

### Supplementary Fig. 1

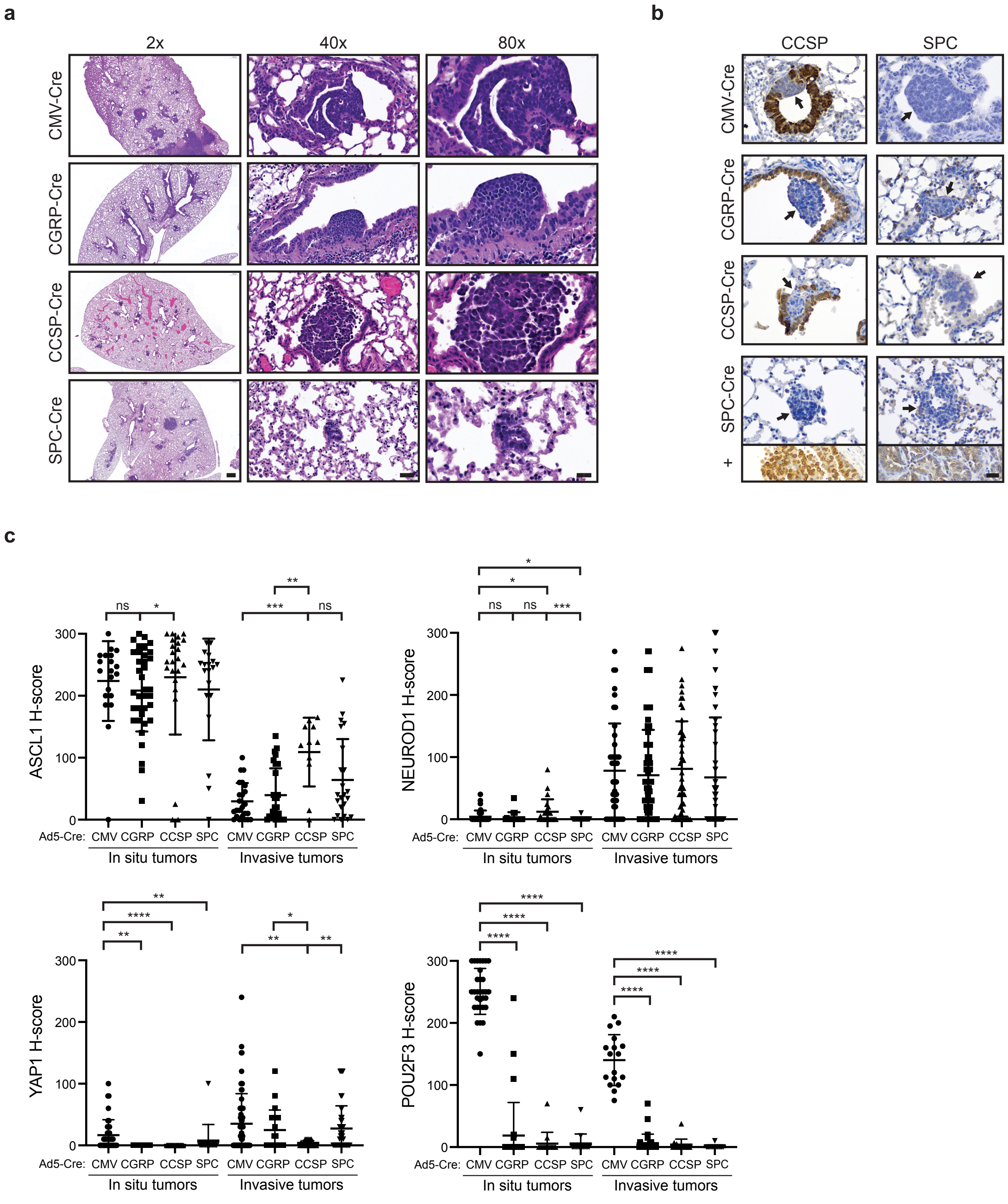

#### Supplementary Figure Legends

**Supplementary Fig. 1, related to Fig. 1: MYC-driven SCLC can arise in multiple lung cell types and initially expresses ASCL1.**

a) Representative H&E images of early-stage RPM tumors initiated with the indicated cell type-specific Cre viruses. Scale bars represent 500 (2x), 40 (40x) and 20 (80x) µm, respectively.

b) Representative IHC images for CCSP and SPC antibodies (top labels) from in situ tumors from RPM mice infected with cell-type specific Cre viruses. Arrows indicate in situ tumors. Inset in bottom panels represent adenocarcinoma positive controls from *Kras*$^{G12D/+}$;*Trp53*$^{fl/fl}$ (KP) mice. Scale bar: 25 µm.

c) IHC quantification from Fig. 1c and 1d with in situ and invasive tumors categorized by tumor-initiating virus (Ad5-Cre). Approximately 10-85 tumors from 3-10 mice per condition were quantified. Mean ± SD. Mann-Whitney two-tailed t-test. * $p < 0.05$; ** $p < 0.01$; *** $p < 0.001$; **** $p < 0.0001$; ns = not significant.

### Supplementary Fig. 2

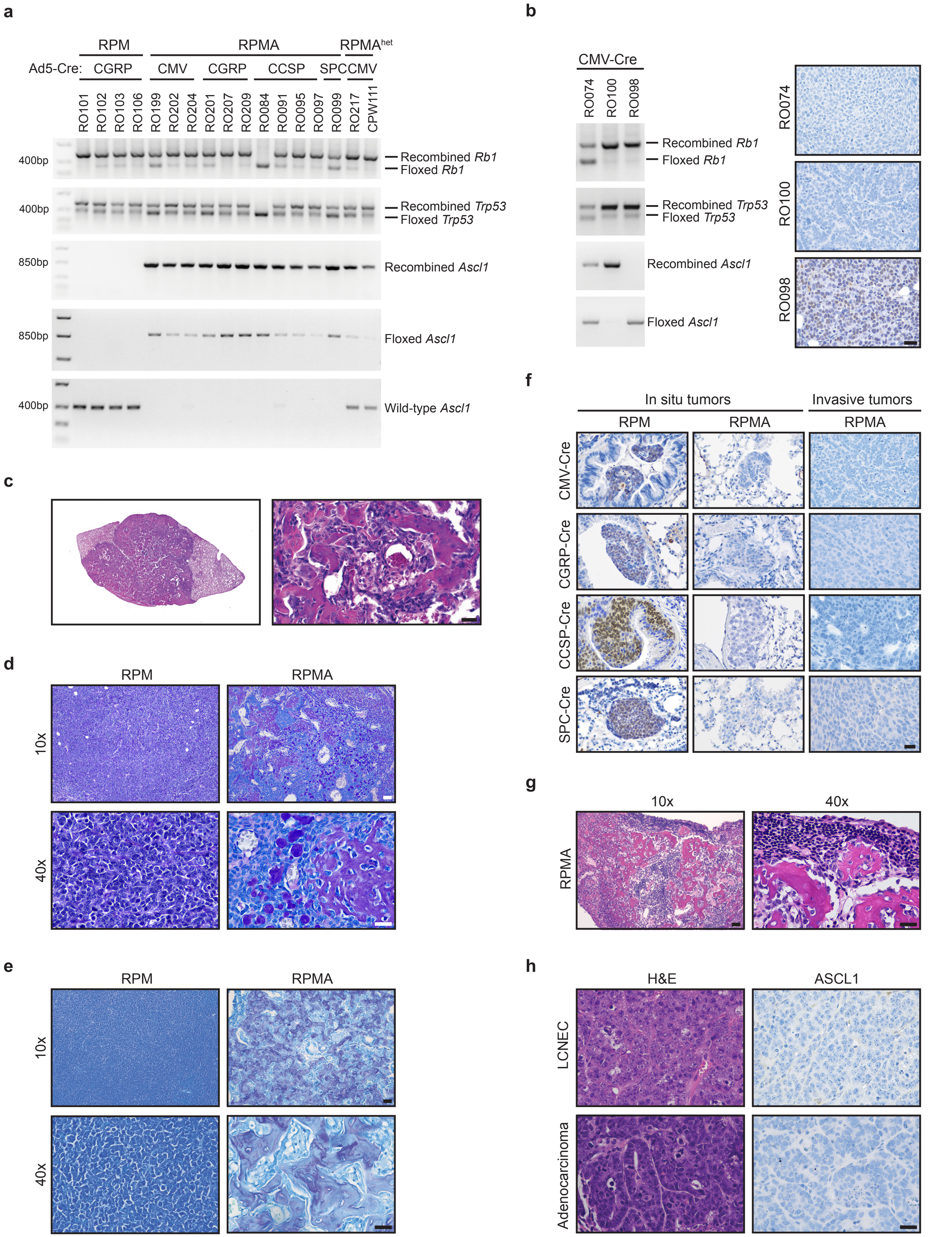

**Supplementary Fig. 2, related to Fig. 2: ASCL1 loss delays tumorigenesis and promotes bone differentiation in multiple cells of origin.**

a) PCR for recombination of the *Rb1*, *Trp53*, and *Ascl1* alleles in individual micro-dissected RPM and RPMA tumors.

b) PCR for recombination of the *Rb1*, *Trp53*, and *Ascl1* alleles in three additional micro-dissected RPMA-CMV tumors, along with corresponding ASCL1 immunohistochemistry. Sample RO098 did not exhibit *Ascl1* recombination and consistently had detectable ASCL1 protein. Scale bar: 25 μm.

c) Representative H&E staining of an RPMA-CGRP osteosarcoma. Scale bar: 25 μm.

d) Representative PAS with Alcian Blue staining (PAB) in RPM-CMV vs RPMA-CMV tumors. Scale bar indicates 50 μm in 10x image (top) and 25 μm in 40x image (bottom).

e) Representative Toluidine Blue staining in RPM-CGRP vs RPMA-CGRP tumors. Scale bar indicates 50 μm in 10x image (top) and 25 μm in 40x image (bottom).

f) Representative IHC for ASCL1 in RPM (positive control) vs early (in situ) RPMA tumors compared to advanced (invasive) RPMA tumors initiated with the indicated cell-type-specific Cre viruses. Scale bar: 25 μm.

g) Apparent osteosarcoma metastasis to the lymph node in an RPMA-CGRP mouse with high lung tumor burden. Scale bar indicates 50 μm in 10x image (left) and 25 μm in 40x image (right).

h) Representative histology and ASCL1 expression in a subset of RPMA tumors with adenocarcinoma or large cell neuroendocrine (LCNEC) morphology. Scale bar: 25 μm.

### Supplementary Fig. 3

**a**

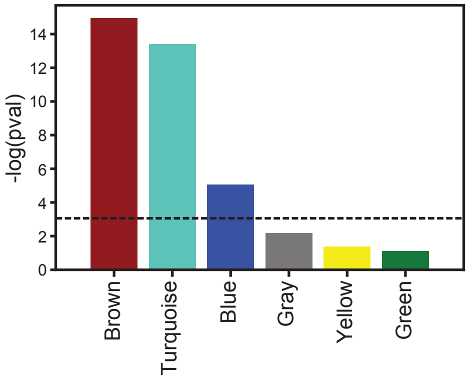


**b**

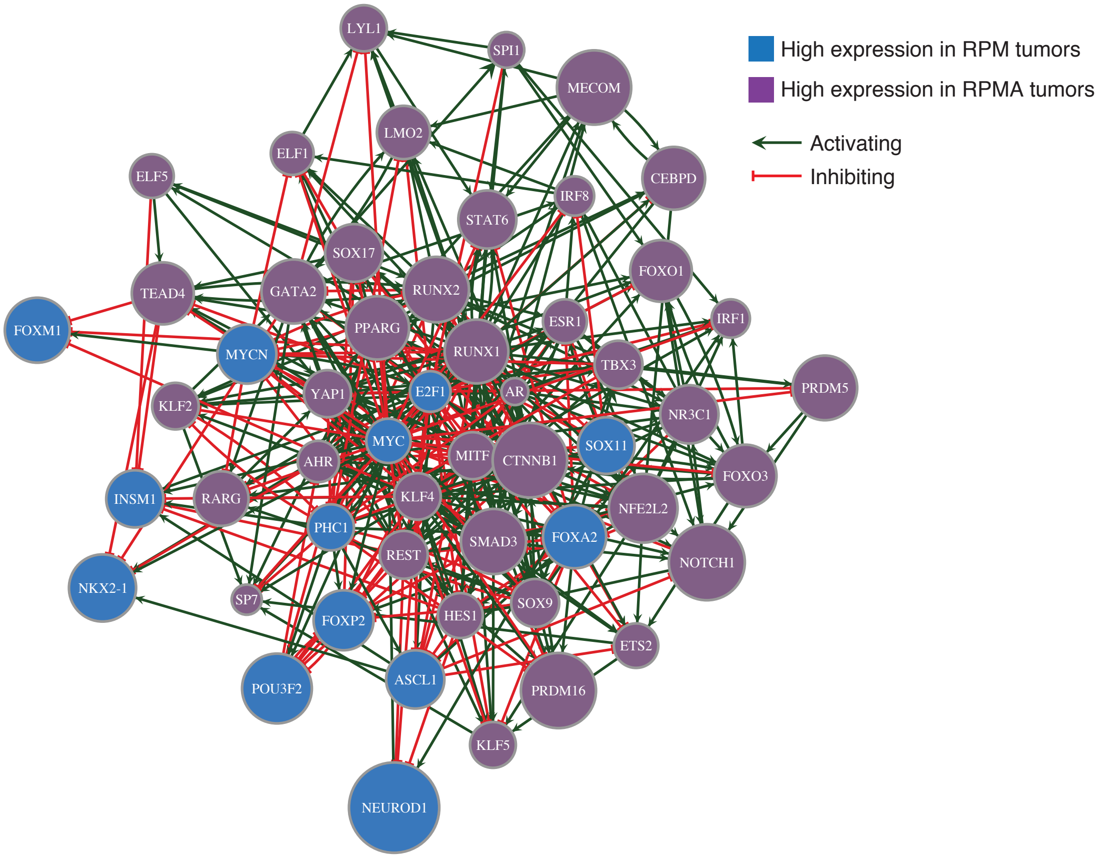

**Supplementary Fig. 3, related to Fig. 4: Network analyses predict transcriptional regulators that drive osteosarcoma cell fate upon ASCL1 loss**

a) ANOVA statistical analysis of co-regulated gene modules identified by Weighted Gene Coexpression Network Analysis (WGCNA) in RPM (n=11) vs RPMA (n=6) tumors. Three modules had significant differential gene expression. The turquoise module is highly expressed in RPMA tumors, and brown and blue modules are highly expressed in RPM tumors. Data is shown as negative log10-transformed p-value. Dotted line indicates p = 0.05.

b) Predicted transcription factor interaction network from RPM vs RPMA tumors based on the genes most central to each identified differentially expressed gene module and data from ChIP-seq databases.

### Supplementary Fig. 4

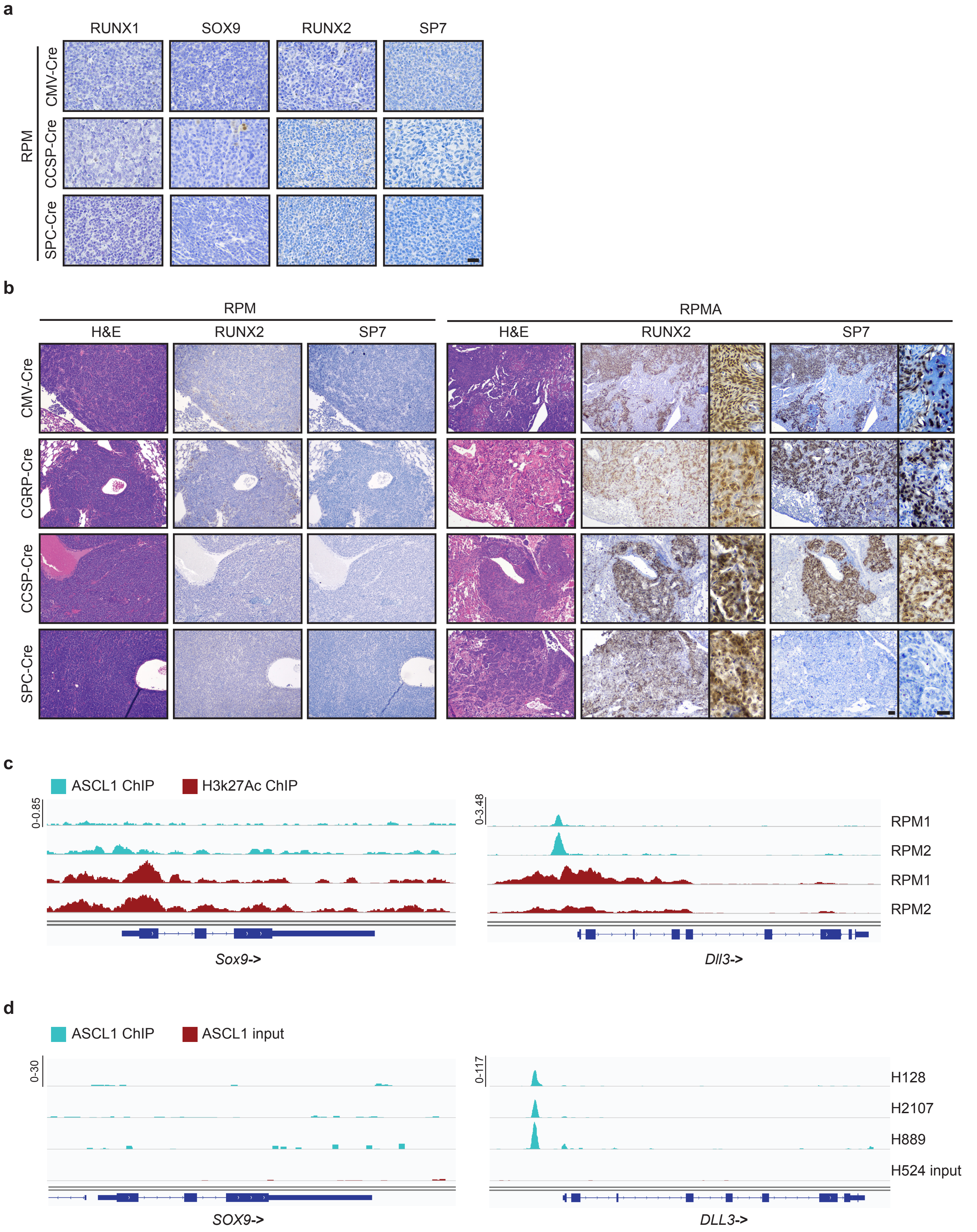

**Supplementary Fig. 4, related to Fig. 6: ASCL1 represses SOX9 and non-endodermal cell fates.**

a) Representative IHC for RUNX1, SOX9, RUNX2 and SP7 in RPM tumors initiated with the indicated Cre viruses. Scale bar: 25 µm.

b) Representative serial sections of RPM or RPMA tumors from mice infected with cell-type-specific viruses stained with H&E, RUNX2, and SP7 antibodies. Scale bar: 50 µm; high magnification inset scale bar in RPMA panel: 25 µm.

c) ChIP-seq analysis of ASCL1 (light blue) and H3K27Ac (red) genomic binding at indicated gene loci from n = 2 independent RPM tumor samples labeled on right panels. Blue rectangles below plots indicate gene exons with directionality of gene (->) near gene name. *Dll3* serves as positive control.

d) ChIP-seq analysis of ASCL1 (light blue) and input control (red) genomic binding at indicated gene loci from human SCLC cell lines labeled on right panels. Blue rectangles below plots indicate gene exons with directionality of gene (->) near gene name. *DLL3* serves as positive control.

**Supplementary Videos:** Movies depicting microCT bone analysis from RPM and RPMA mice infected with either CMV-Cre or CGRP-Cre as indicated. Inset image is axial cross-section showing tumor burden by microCT.

**Supplementary Table 1:** Differentially expressed genes obtained from RNA-seq analysis, curated gene lists used for GSEA, and Panther and REViGO outputs for RPMA vs RPM tumors.

**Supplementary Table 2:** Data from IPA comparing gene expression and predicted upstream regulators in RPMA vs RPM tumors.
