## Supplementary figures and images for "ASCL1 represses a latent osteogenic program in small cell lung cancer in multiple cells of origin"

### Supplementary Videos

#### Slide 1
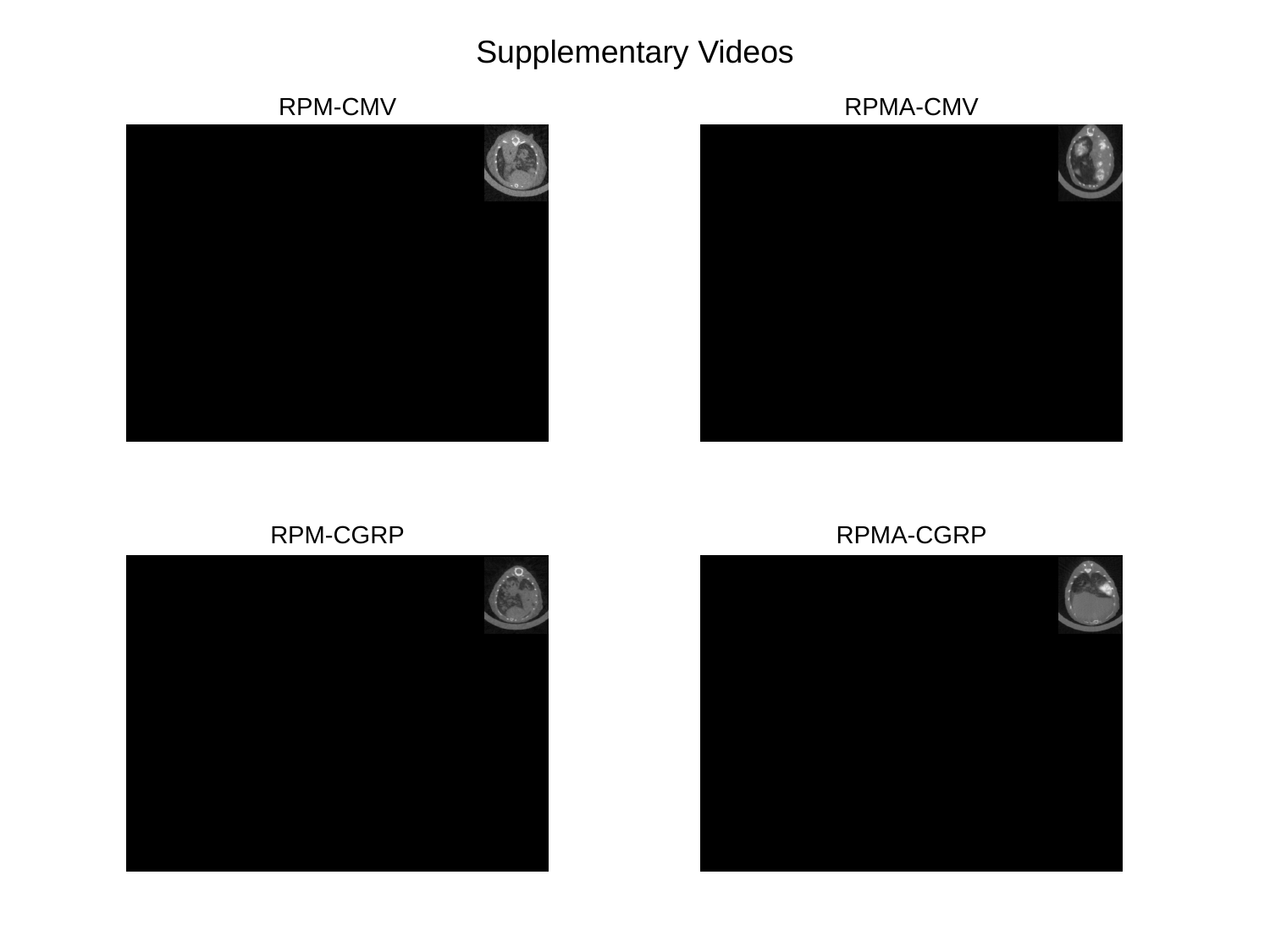

Supplementary Videos
RPM-CMV
RPMA-CMV
RPM-CGRP
RPMA-CGRP
